# Hepatic management of toxic sterols after acute deletion of *Cyp51* from cholesterol synthesis

**DOI:** 10.1101/2025.09.18.674867

**Authors:** Tinkara Kreft, Kaja Blagotinšek Cokan, Cene Skubic, Tadeja Režen, Martina Perše, Karmen Wechtersbach, Nika Kojc, Jera Jeruc, Željko Debeljak, Marija Heffer, Madlen Matz-Soja, Damjana Rozman

## Abstract

Lanosterol 14α-demethylase (CYP51) is an enzyme involved in cholesterol synthesis, crucial for the normal liver function. Diminished activity of CYP51 leads to metabolism associated liver disease, ending in hepatocellular carcinoma. It is still not clear which processes are most affected in the hepatocytes and how they communicate with other cells towards the progressive liver pathology. Herein we describe a new inducible, liver-specific *Cyp51* knockout mouse model (iLKO), developed to study how acute disruption of cholesterol synthesis is managed in the adult liver. Doxycycline inducible deletion avoided developmental confounders of albumin-Cre models and enabled isolation of viable primary hepatocytes. iLKO hepatocytes and liver tissue showed CYP51 depletion with accumulation of lanosterol and 24,25-dihydrolanosterol, while hepatic cholesterol levels remained largely unchanged, indicating compensatory uptake and/or pathway rerouting. Histological examination and transmission electron microscopy (TEM) revealed hepatomegaly with mild portal inflammation and ductular reaction but no overt fibrosis at the studied time points. iLKO hepatocytes displayed increased nuclear lipid droplets (LD) that might be involved in adaptation to endoplasmic reticulum (ER) stress. Additionally, we discovered crystal-like inclusions, especially in Kupffer cells. MALDI-TOF MSI could not resolve their composition, but their occurrence alongside sterol overload implicates non-cholesterol sterol crystallization as a potential trigger of inflammation. In summary, iLKO is a suitable model for dissection of sterol toxicity, clearly separated from developmental effects of *Cyp51* depletion. Future examinations could reveal how toxic sterol intermediates are buffered in adult liver and how they might be connected to inflammation driven pathologies, relevant also to human health.

## 1. Introduction

Cholesterol biosynthesis is a tightly regulated, multistep metabolic pathway where each enzymatic reaction not only contributes to the formation of cholesterol but also regulates the levels of diverse sterol intermediates. Many of which have emerging, yet still poorly understood, biological functions [1,2]. Among the key enzymes in this pathway is lanosterol 14α-demethylase (CYP51), which catalyzes the first step in the post-squalene segment. CYP51 is highly expressed in the liver, a primary site of *de novo* cholesterol synthesis and systemic sterol regulation [3,4].

The essential role of CYP51 in mammalian development and physiology has been explored in vivo through a series of *Cyp51* knockout (KO) models. The earliest, developed by Keber et al. [5], used a global *Cyp51* KO, uncovering *Cyp51* essential function during embryogenesis, as KO mice died mid-gestation due to cardiovascular malformations. Based on a comparative analysis of developmental defects, the authors noted a strong phenotypic resemblance to Antley-Bixler syndrome in humans.

To bypass embryonic lethality of the first mouse model, Lewinska et al. [6] analyzed heterozygous *Cyp51 KO (Cyp51^+/−^)* mice. While these animals appeared grossly normal, deeper analysis revealed sex-and diet-dependent metabolic alterations, including elevated cholesterol levels, increased hepatic apoptosis, and compensatory gene expression changes, especially under a high-fat, high-cholesterol diet. Interestingly, males displayed more pronounced hepatic stress, whereas females showed a more adaptive gene regulatory response. These findings demonstrated that even partial loss of CYP51 can lead to subclinical pathologies, highlighting its dosage sensitivity and interactions with environmental factors like diet and sex.

To enable studies of liver-specific functions, Lorbek et al. [7] established a hepatocyte-specific *Cyp51* KO model (LKO), using Cre-LoxP system with Cre recombinase expressed under the albumin *(Alb)* promoter. They characterized the effects of *Cyp51* deletion in young adult LKO mice, uncovering early signs of fibrosis, oval cell activation, and sex-specific transcriptional reprogramming, including down-regulation of primary metabolism pathways in females and a stronger immune response in males. The LKO model subsequently served as the foundation for two temporally distinct studies. Urlep et al. [4] examined this model during neonatal liver development, revealing that early CYP51 loss disrupts hepatocyte differentiation, triggers endoplasmic reticulum (ER) stress, and induces a ductular reaction. Cokan et al. [8] extended these observations in aged LKO mice, hypothesizing that long-term sterol imbalance leads to disease progression. Indeed, aged *Cyp51 KO* mice developed hepatocellular carcinoma (HCC), with higher prevalence in females and sex-specific reprogramming of inflammatory, metabolic, and proliferative gene networks.

In addition to the liver-specific models, Keber et al. [9] created a male germ cell-specific *Cyp51* KO using Stra8-Cre^+^ transgenic mice. Surprisingly, despite efficient deletion, these mice maintained normal spermatogenesis and fertility, suggesting that CYP51 is dispensable in male germ cell development, highlighting tissue-specific requirements for sterol biosynthesis. More recently, Skubic et al. [10] also developed a cell-based *CYP51* KO model using HepG2 human hepatoblastoma cell line, allowing in-depth analysis of sterol-related mechanisms in a human hepatic context that mirrors features observed in murine liver-specific KOs.

Despite the valuable insights provided by these models, the current liver-specific model LKO has its own limitations. The major one lies in the timing of gene deletion driven by the albumin promoter. The Alb-Cre system becomes active during fetal liver development, beginning at ED 10.5 [11,12], which results in *Cyp51* recombination prior to full hepatic maturation. This early onset complicates the interpretation of downstream phenotypes, as observed changes may originate from disrupted liver development rather than from adult hepatocyte-specific metabolic defects. Another critical drawback is the model’s unsuitability for primary hepatocyte isolation. By the time the liver phenotype becomes fully established, typically around 19 weeks of age, mice exhibit pronounced fibrosis, inflammation, and ductular reactions [7]. These pathological changes compromise the yield and viability of isolated hepatocytes, limiting the use of this model for in vitro studies of sterol metabolism.

To overcome these limitations, we developed an inducible liver-specific *Cyp51* knockout model (iLKO), which allows controlled induction of CYP51 loss in adult mice with fully developed liver. Our aim was to characterize this model and assess the effects of knockout. Importantly, the system also enables the isolation of viable primary hepatocytes from phenotypically relevant but histologically intact livers, facilitating direct mechanistic studies.

## 2. Materials and Methods

### 2.1. Transgenic iLKO mouse model

The iLKO mouse model was obtained by crossbreeding the rTA^LAP^-1/LC1-1 mice [76] and the Cyp51^flox/flox^ mice (B6.129SV*-Cyp51*^tm1Bfro^; *MGI code: 4357830) [5] (***Figure 1**). The rTA^LAP^-1/LC1-1 mice transgenic mice were created by prof. dr. Rolf Gebhardt group at the Institute of Biochemistry, Faculty of Medicine, University of Leipzig, Germany to enable the tissue- and time-specific gene deletions in hepatocytes. The resulting triple transgenic mice of F1 generation were then intercrossed to obtain iLKO mice carrying (1) two *Cyp51* alleles with loxP sites flanking exons 3 and 4, (2) the reverse tetracycline-controlled transactivator *(rtTA) [77]* under the liver-specific P_LAP_ promoter [78], and (3) the Cre recombinase gen *(Cre)* under tetracycline-responsive control. Knockout was induced with administration of a tetracycline analogue, doxycycline *(Dox)*, via drinking water. This antibiotic triggers transcription of *Cre through rtTA*, which is constitutively expressed in hepatocytes. Cre recombinase cuts out a sequence between two *loxP* sites, leading to loss of functional CYP51 in hepatocytes. Control mice (WT) were littermates obtained from the same parental crosses, but identified by genotyping as lacking the *Cre* transgene, and therefore did not undergo *Cyp51* deletion following doxycycline treatment.

**Figure 1.**
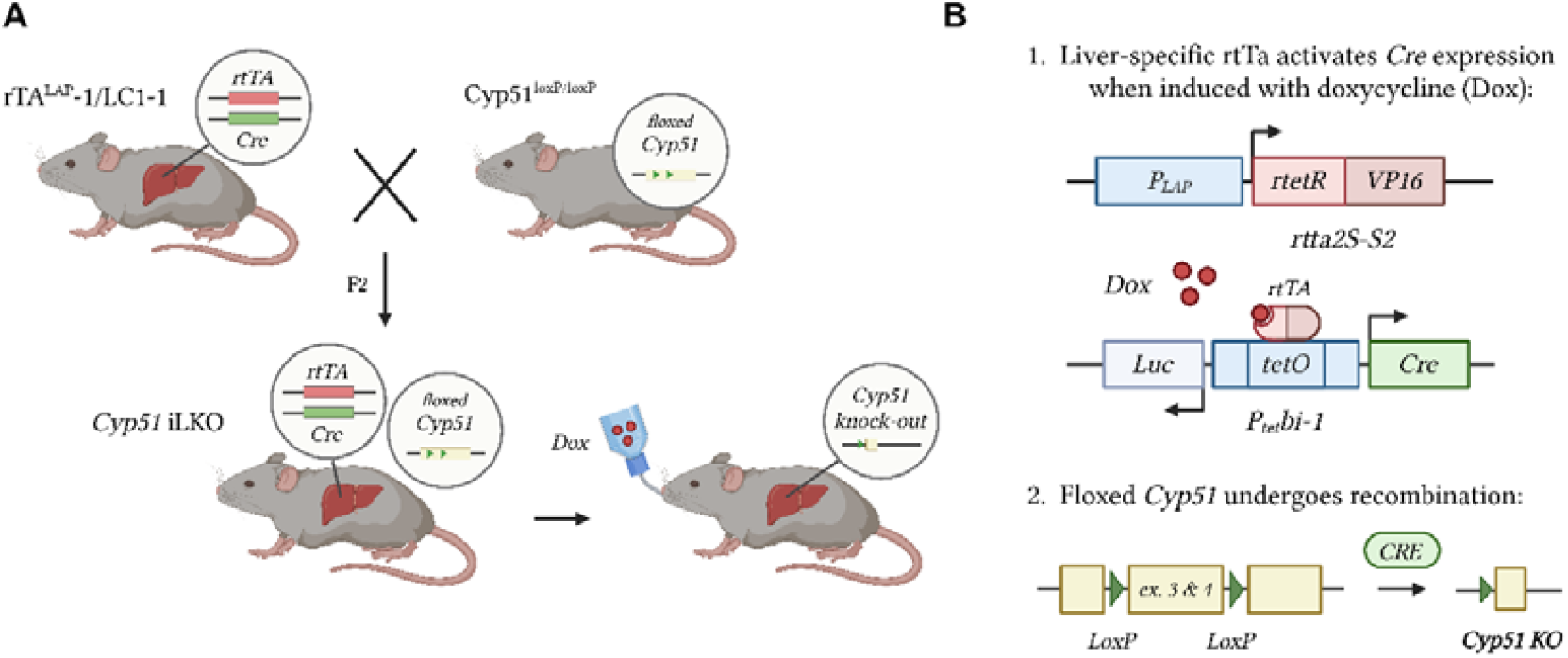
Generation of transgenic iLKO mouse model **(A)** Crossbreeding rTALAP-1/LC1-1 mice with *Cyp51*^flox/flox^ mice produces triple transgenic F2 offspring carrying two floxed *Cyp51* alleles, the reverse tetracycline-controlled transactivator (rtTA) and the Cre recombinase gen (Cre) under tetracycline-responsive control. Deletion of *Cyp51* is induced by administering doxycycline (Dox) in the drinking water. **(B)** Schematic representation of Cyp51 deletion process. First, the liver-specific P_LAP_ promoter drives expression of a fusion protein rtTA, composed of a tetracycline-sensitive rtetR domain and the VP16 transcriptional activator. In the presence of Dox, rtTA binds to the tetracycline operator sequences (tetO) located within a bidirectional promoter P_tet_bi-1, thereby activating the expression of both Cre recombinase and luciferase (*Luc*). Second, Cre mediates site-specific recombination between LoxP sites flanking exons 3 and 4 of the *Cyp51* gene, resulting in its deletion. Figure was created with BioRender.

Genotypes were identified by PCR using DNA from ear biopsies of 4-5 weeks old mice. For PCR analyses, the following primers were designed as seen in **Table S1**. PCR resulting material was analyzed by agarose gel (1-1.5%) electrophoresis (**Figure S1**).

### 2.2. In vivo animal procedure and ethics statement

Animals were bred under the standard condition for laboratory animals: 12 h light/dark cycle (7:00 am - 7:00 pm light), 55±10%, humidity and at 22±1°C, using standard rodent chow (diet 1324, Altromin, Germany) and acidified tap water (pH = 3) *ad libitum*. They were housed in groups of 3-5 mice per cage. The results of negatively tested microorganisms of routine health microbiological monitoring are shown in **Table S2**.

Mice aged 9-10 weeks were administrated 2 mg/ml Doxycycline Hyclate (Sigma-Aldrich, St. Louis, MO, USA, St.Louis, MO; Dox) in drinking water for 10 days. The Dox injections were repeated 3-times at intervals of 3-4 days. During Dox application, all mice were weighed and clinical examinated every day till euthanasia.

Mice were sacrificed a day after last Dox application at age 10-11 weeks (in text: 1d post-Dox) and 14-15 weeks (in text: 4w post-Dox) with cervical dislocation between 11-13h (to minimize diurnal variations and after a 4-6 h of fast (food withdrawal at 7 a.m.)) or under isoflurane anaesthesia (Isoflurin, 1000mg/g stock solution; Vetpharma Animal Health, Spain) for primary hepatocytes isolations (without food withdrawal before the procedure and in the period between 10-13 h).

After cervical dislocation the blood was taken immediately after cervical dislocation from the right ventricle, collected into heparin-coated Vacuette MiniCollect® 1 ml Plasma Tubes (Greiner Bioone, Frickenhausen, Germany) and centrifuged for 4°C, 3000 g for 15 min and the organs (liver, kidney, spleen, heart, gonads, stomach, and intestine) removed and patho-morphologically examined. The left lateral liver lobus was taken for histology, part of the right lateral liver lobus was prepared for TEM and other parts were cut into thin slices, snap-frozen in liquid nitrogen and stored at – 80°C for subsequent analysis. For this research, only livers were used.

Mouse experiment was approved by the Veterinary Administration of the Republic of Slovenia (license number U34401-10/2016/5) and was conducted in agreement with the National Institute of Health guidelines for work with laboratory animals, national legislation, and Directive 2010/63/EU on the protection of animals used for scientific purposes.

### 2.3. Histological liver examination

The formalin-fixed paraffin-embedded livers were sectioned onto 4-5 µm glass slides and stained with H&E for standard histological analyses. Briefly, liver sections were deparaffinized at 70°C for 10 min and washed in two changes of xylene, 100% ethanol, 95% ethanol, 70% ethanol, and twice in acidified water. Liver sections from 70 mice were examined under the light microscope for hepatocyte morphology, portal tract abnormalities (ductular reaction) and the presence and localization of inflammatory infiltration and lipid droplets. The intensity of inflammation and the ductular reaction was graded as normal (0), mild (1), moderate (2), and severe (3). We analyzed H&E data using three-way ANOVA. For more concise visualization of the main effects, pooled-sex data were additionally analyzed using two-way ANOVA with Fisher’s test.

### 2.4. HepG2 cell culture

HepG2 cell line, a human male liver cancer cell line, (Synthego, MenaloPark, CA, USA) with CRISPR-Cas9 mediated *CYP51* deletion and the native HepG2 cell line, were obtained from Synthego (Synthego, Menalo Park, CA, USA) with a certificate of authentication and a negative mycoplasma test. Both have been previously described and validated for *CYP51* expression by our group [10]. Cells were maintained in Dulbecco’s Modified Eagle’s Media (DMEM) high glucose, supplemented with 10% fetal bovine serum (FBS) and 1% penicillin-streptomycin (P/S), all from Sigma-Aldrich, in a 5% CO_2_ incubator at 37 °C.

For transmission electron microscopy (TEM) analysis of crystal formation, cells were seeded onto 100 mm plates at a density of 2 × 10^6^ cells/plate and kept for 24 h in normal culturing conditions described above. After 24 h, culture medium was removed, cells were washed twice with phosphate buffered saline (PBS), and lipid-depleted medium was added (DMEM high glucose supplemented with 10% lipid-depleted serum (Biosera inc., France), 1% P/S and 30 mg/ml of cholesterol (Sigma Aldrich) dissolved in ethanol). After 48 h in lipid-depleted medium, cells were washed with PBS, scraped free in PBS and then collected to fresh tube. Following centrifugation (1000 g, 5 min), cell pelet was resuspended in solution for fixation, followed by the steps described in section 2.5. *Transmission electron microscopy (TEM)*.

### 2.5. Transmission electron microscopy (TEM)

Mouse liver tissue samples and HepG2 cell pellet samples were fixed in a solution of neutral buffered 10% formaldehyde for 1h, washed with 0.1 M Millonig’s phosphate buffer, postfixed in a solution of 1% osmium tetroxide (Merck, Darmstadt, Germany) for 30 min and washed with distilled water. HepG2 cell pellet samples were additionally embedded in HistoGel™ (Epredia, Kalamazoo, MI, USA). Both HepG2 and tissue samples were then dehydrated in increasing concentrations of ethanol and later in 1-2-propylene oxide (Merck, Darmstadt, Germany) for 10 min each. Samples were then incubated in a mixture (1:1) of 1-2-propylene oxide and Epon 812 resin (Serva Electrophoresis, Heidelberg, Germany) for 20 min, embedded in 100% Epon, and polymerized overnight at 60 °C. Semithin sections (1.4 µm) were cut using a Leica EM UC6 ultramicrotome (Leica Microsystems, Wetzlar, Germany), stained with Azur II staining solution (Merck, Darmstadt, Germany), and analyzed by light microscopy. Ultrathin sections (60 nm) were counterstained with 5% uranyl acetate (SPI Supplies, West Chester, PA, USA) and 3% lead citrate (Merck, Darmstadt, Germany) and examined by 120-kV JEM-1400 transmission electron microscope (JEOL, Tokyo, Japan).

Sample of each mouse liver was scored for size and shape of the nucleus, mitochondria, and ER of hepatocytes, proliferation of bile ducts, presence of portal and parenchymal inflammation, and assigned as NO (absence of ductular proliferation or inflammation) or YES (presence ductular proliferation or inflammation). The incidence of lipid droplets in hepatocytes and crystals in hepatocytes and Kupffer cells was graded as: normal (0), mild (1), moderate (2), and severe (3). TEM data were analyzed using three-way ANOVA. To provide more concise visualization of the main effects, pooled-sex data were additionally analyzed using two-way ANOVA with Fisher’s test.

### 2.6. Primary hepatocytes isolation with two-step collagenase perfusion technique

Before the isolating hepatocytes procedure, all necessary reagents and buffers (listed in *Table S3*) were prepared. The cell culture plates (TPP, Switzerland) were pre-coated with 0.01% rat-tail collagen (self-made; stock solution: 100%, 1 mg/ml), incubated at 37 °C under the sterile condition in a humidified CO_2_ incubator for 12h, washed once with Hanks^+/+^ + EDTA and twice with Hanks^+/+^ and stored in Hanks^+/+^ in the fridge at 4 °C.

The perfusion buffer (PPM) was placed into a sterilized, washed and prewarmed peristaltic pump tubing (to 40 °C). The tubing was primed with warmed PPM (perfusion pump speed was 8-10 ml/min) and avoided all bubbles through the system.

Following 1.5-3% isoflurane anesthesia, the mouse was positioned on the dissection tray. We wet the fur thoroughly with 70% ethanol and made an incision through the skin that was secured near the head using a needle. After the stomach and intestinal removal from the body, the portal vein was cannulated, inferior vena cava was cut and the liver was perfused with PPM to wash out blood. Further, another needle was inserted through the right part of the heart into superior vena cava and the inferior vena cava was closed. After the preparative step for liver digestion, we perfused the liver with Collagenase, glucose and CaCl_2_ solution in the prewarmed and oxygenized perfusion buffer (PM) to the liver to digest collagen in the extracellular matrix for 6-8 min. Using forceps, the liver was dissected out gently by cutting all the connections of the liver to other organs and the gall bladder was removed. The liver was placed in the falcon tube with cold isolation medium (IMS), where cells were released into suspension. Viable hepatocytes were separated from dead and non-hepatocyte cells by three centrifugations (1x 2 min, 600 rpm; 2x 5 min, 500 rpm), replacing the supernatant with fresh IMS between steps.

### 2.7. Culturing of primary mouse hepatocytes

After isolation, primary hepatocytes of one mouse were released into cold IMS buffer, centrifuged 3-times at 4 °C (once at 600 rpm for 2 min and twice at 500 rpm for 5 min) and the supernatant was aspirated. Cell viability and count were assessed using a 1:1 dilution of Trypan blue (Sigma-Aldrich, St. Louis, MO, USA). Cells were then plated in triplicates in 6-well (2.5-3 × 10^5^ cells/well) and quadruplicates in 12-well (1-1.5 × 10^5^ cells/well) collagen pre-coated plates containing attachment medium (William’s E medium (Gibco), 2mM L-glutamine (Sigma-Aldrich), 1% P/S solution (penicillin: 100 units/ml and streptomycin: 0.1 mg/ml; Sigma-Aldrich) and 100 nM dexamethasone (Sigma-Aldrich)) in a humidified 5% CO_2_ incubator at 37 °C. After 3 h, we changed the medium with a cultivation medium (attachment medium with 10% FBS (Sigma-Aldrich)). After 24 h of plating, primary hepatocytes acquired their typical hexagonal shape as seen in **Figure S2**. We harvested cells with resuspended in PBS twice, aspirated supernatant and the pellets were freshly frozen at -80 °C for further RT-qPCR, western blot, and sterol analyses.

### 2.8. Western blotting

Total proteins were isolated from primary hepatocytes using RIPA lysis buffer (20 mM Tris/HCl pH 8, 150 mM NaCl, 1% NP-40, 0.5% sodium deoxycholate, 0.1% SDS) containing 1 mM PMSF and cOmplete protease inhibitor cocktail (Roche, Basel, Switzerland). Cell pellets were diluted in 200 µL buffer and sonicated for 5–10 min at max using Bioruptor UCD-200 (Diagenode, Liège, Belgium). After centrifugation at 12000g, 15 min at 4 °C, we measured protein concentration using Pierce™ BCA Protein Assay Kit (Thermo Fischer Scientific, Waltham, MA, USA) according to the manufacturer’s instructions. Western blot was performed according to Abcam’s instructions. 5 µg of proteins mixed with NuPAGE® LDS Sample Buffer (Thermo Fischer Scientific, Waltham, MA, USA) in final volume 20 µL and 5 µl of PageRuler Plus Prestained protein ladder (Thermo Fisher Scientific, Waltham, Massachusetts, United States) were loaded. Samples were separated on a 10% SDS-PAGE gels and transferred on Immobilon-P PVDF membranes (Merck KGaA, Darmstadt, Germany) using Mini-PROTEAN System (Bio-Rad Laboratories, Hercules, California, United States).

Anti-CYP51 antibodies (1:2000; custom-made [5] rabbit polyclonal antibody against peptide QRLKDSWAERLDFNPDRY) and HRP-conjugated goat anti-rabbit IgG H&L secondary antibodies (1:20.000; Abcam, cat.no. ab205718) were used. For loading control, mouse monoclonal anti-GAPDH antibodies (1:10.000; Abcam, cat.no. ab8245) and HRP-conjugated goat anti-mouse IgG secondary antibodies (1:20.000; Santa Cruz, cat.no. sc-2005) were applied. Visualization of CYP51 and GAPDH was done with SuperSignal™ West Femto (Thermo Fisher) and Immobilon® Classico Western HRP Substrat (Merck), respectively and chemiluminescence was recorded using a Fujifilm LAS 400 system (Fujifilm, Singapore). ImageJ was applied to analyze images. We used a Mann-Whitney test and a one-tailed p-value threshold of 0.05 as a measurement of significance. Results are presented as mean ± SEM.

### 2.9. RNA isolation and reverse transcription-quantitative PCR (RT-qPCR)

The primary hepatocyte samples we applied for total RNA isolation using TRI Reagent (Sigma-Aldrich, St. Louis, MO, USA, St. Louis, MO, USA) procedure. RNA concentration and purity of frozen samples were assayed using NanoDrop 1000 Spectrophotometer (Thermo Fischer Scientific, Waltham, MA, USA). To quantify the genes of interest expression levels, equal amounts of cDNA were synthesis with amplification grade DNase I (Roche, Basel, Switzerland), and Transcriptor Universal cDNA Master (Roche, Basel, Switzerland) from initial 2 µg RNA of samples according to manufacturer’s instructions. *Actb, Utp6*, and *Hmbs* were used as the internal controls for each sample.

For the real-time quantitative reverse transcription polymerase chain reactions (RT-qPCR) we used PCR reaction consisted of 2.5 µL SYBR Green I Master (Roche, Basel, Switzerland), 0.75 µL of cDNA template, 0.6 µL of 2.5 µM random primer mix and 1.15 µL of RNAse-free water in a final volume of 5 µL. All performed qPCR using SYBR Green was conducted at 10 min incubation at 95 °C followed by 45 cycles of 10 s at 95 °C, 20 s at 58 °C and 20 s at 72 °C. Experiments were made on a Roche LightCycler 480 (Roche, Basel, Switzerland) in three technical replicates for each of the sample using 384-well plates. The specificity of the reaction was verified by melt curve analysis.

Relative transcript levels were calculated by the comparative Cp method and -ΔΔCp values were used for statistical analyses. We used Mann-Whitney test and a p-value threshold of 0.05 for measurement of significance. The primer sequences in RT-qPCR are listed in *Table S4*.

### 2.10. Sterol extraction and liquid chromatography with tandem mass spectrometry (LC-MS/MS)

Sterols were extracted from primary hepatocytes and frozen liver tissue samples according to the previously described protocol [79]. Briefly, frozen cell pellets were resuspended in Folsch reagent (chloroform/methanol; 2:1), vortexed and then 200 ng of internal standard lathosterol-D7 (Avanti Polar Lipids) was added. Samples were incubated for 2 h at room temperature, with shaking. In case of tissue samples, 1 ml of Folsch reagent was added to 1 mg of ground frozen liver tissue and incubation was overnight. After incubation, Folsch reagent was evaporated from the extract and incubation in hydrolysis solution (8 g NaOH + 20 ml H_2_O + 180 ml 99.5 % ethanol) followed for 1 h at 65 °C. With the addition of cyclohexane two repeated extractions were then made. Cyclohexane from final samples was evaporated and samples were dissolved in methanol for analysis.

The hepatocyte samples were analyzed on the LC-MS/MS according to the previously described protocol by Skubic et al. [80]. The chromatographic separation of sterols was performed on combined columns Luna® 3 µm PFP (2) 100 Å (100 mm and 150 mm length) on Shimadzu Nexera XR HPLC (Shimadzu, Kyoto, Japan). For the mobile phase, 80% methanol, 10% water, 10% 1-propanol, and 0.05% formic acid were used in the isocratic condition. The detection was applied on HPLC, coupled with the Sciex Triple Quad 3500 mass spectrometer. For ionization, APCI (atmospheric-pressure chemical ionization) was used in positive mode and detection was made in MRM (multiple reaction monitoring) mode. Sterols in liver tissue were measured using a slightly adapted protocol [10]. All post-squalene sterol concentrations were calculated to corresponding standards from Avanti Lipids. We normalized those concentrations to 100 µg of total proteins and internal standard (Lathosterol-d7) in case of primary hepatocytes and on wet tissue mass and same internal standard in case of liver tissue samples. In the procedure were measured the sterol intermediates displayed in *Table S5*. Two-tailed Mann-Whitney test and a p-value threshold of 0.05 was used as a measurement of significance.

### 2.11. Matrix-assisted laser desorption ionization–time of flight mass spectrometry imaging (MALDI-TOF MSI)

Frozen liver samples from female iLKO and WT mice were cut using cryocutter (*-20°C)* to 5 µm slices, which were then mounted on a metal plate. DHB matrix was deposited on samples by 30 min sublimation and iMScope device was used for the MSI. Instrumental conditions included positive ionization, maximal voltage and D2 LASER diameter (app. 25 µm). LASER intensity and frequency were 45% and 100 Hz, respectively. m/z range was 300 – 800. 306 pixels and 100 spectra per pixel were recorded. Each sample was analyzed in duplicate.

## 3. Results

### 3.1. The inducible hepatocyte knockout mouse model iLKO

The developed iLKO mouse model allows a controlled time-dependent deletion of *Cyp51* in hepatocytes. This is provided by doxycycline (Dox) controlled liver-specific transactivator (rtTA) which controls the CRE-loxP system that exhibits site-specific partial deletion of *Cyp51* (explained in detailed in *2.1. Transgenic iLKO mouse model*).

The design of experiment is shown in **Figure S3**. In brief, at the age of 9 weeks, the iLKO mice and wild-type (WT) mice received Dox in drinking water for 10 days to induce recombination with the CRE-loxP system. One half of the subjects was euthanized one day after last Dox application (at ∼10 weeks of age). The second half was euthanized 4 weeks later, to evaluate delayed effects. The WT mice served as a control to distinguish the effects of *Cyp51* deletion from potential off-target effects of Dox treatment. Liver samples were used to validate knockout efficiency and histologically evaluate a new iLKO model. Isolated primary hepatocytes were additionally characterized and analyzed at *Cyp51* mRNA, protein, and functional level.

During 10 days of Dox application, we observed a decrease in male iLKO body weights compared to WTs, with a significant difference already at day 1 (**Figure 2A**). No changes were observed in females. **Figure 2B** shows that while the body weight at termination was reduced only in the iLKO males, the liver/body weight ratios were significantly increased in both sexes, indicating the initiation of hepatomegaly. Interestingly, in females, bigger increase of the ratio was observed in a group of mice that were terminated 4 weeks after the last Dox application, characterizing hepatomegaly as a prolonged or delayed effect of *Cyp51* deletion. Similar trend is showing also in males. An increase in male body weight in the WT group of 4 weeks post-Dox terminated mice compared to 1 day post-Dox might indicate a normal body growth that typically continues longer into the young adulthood of males than of females.

**Figure 2.**
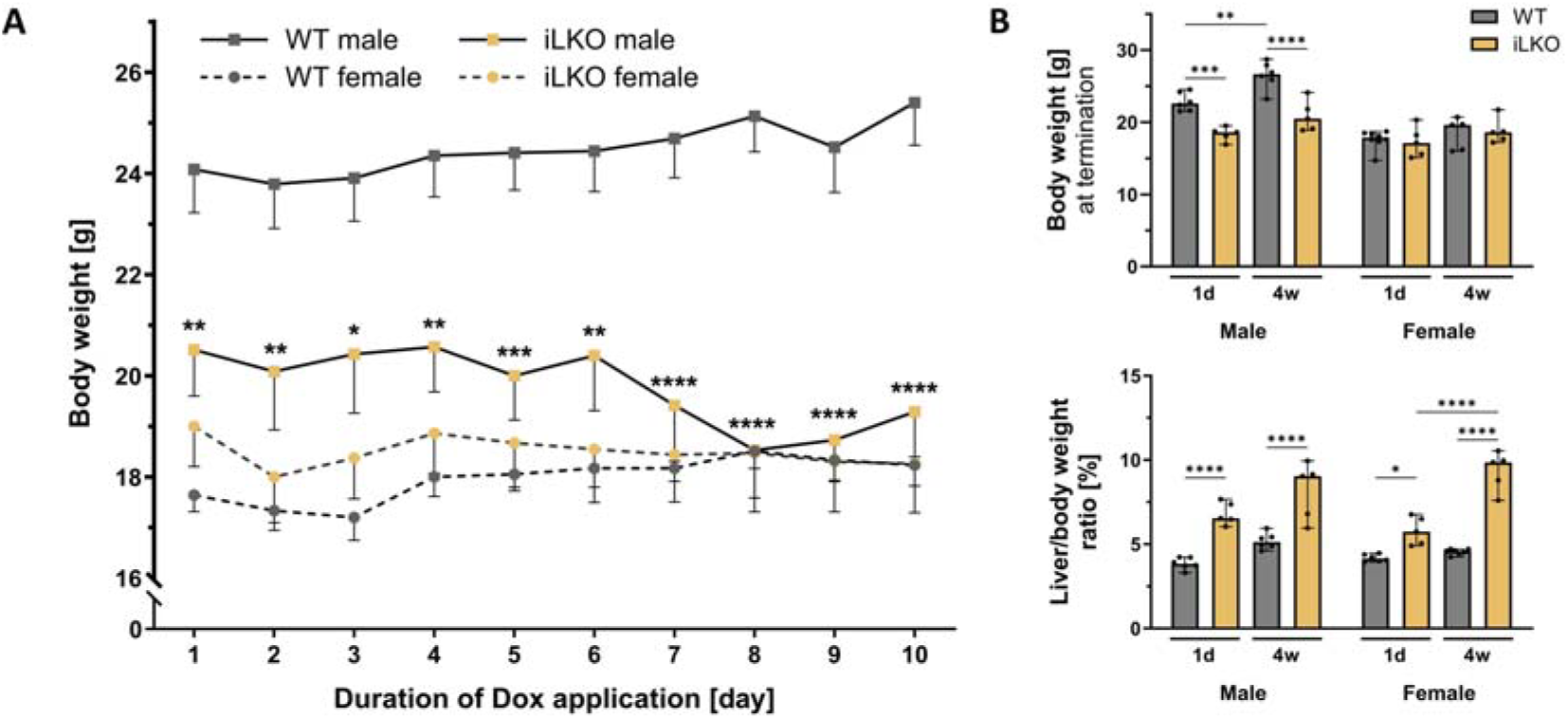
Observations related to body/liver weight. **(A)** Body weight of male and female iLKO and wild type (WT) mice during 10 days of doxycycline (Dox) application (n=7-11/group). Median values with SEM of mice weights were present for each day of Dox application. * p < 0.05; ** p < 0.01; *** p < 0.001; **** p < 0.0001. **(B)** Body weights and liver/body weight ratios of male and female WT and iLKO mice, at termination point (1 day post-Dox or 4 weeks post-Dox) (n=5-8/group). Median values with 95% CI are reported and stat. significance is only shown for WT vs iLKO comparisons. * p < 0.05; ** p < 0.01; *** p < 0.001; **** p < 0.0001.

Seven iLKO mice (six females and one male), that were originally intended to be bred until 14 weeks of age, had to be euthanized prematurely, between 11 and 13 weeks, due to changes in posture, apathetic behavior, and suspected jaundice (**Figure S4A**). Pathoanatomical examination revealed jaundiced mucous membranes, and their gallbladders were filled with bile containing sand-like structures (*Figure S4B, C*). A similar phenotype has been observed earlier in the *Cyp51* LKO model in prepubertal mice [4]. They also observed a higher frequency of severe changes in females.

### 3.2. CYP51 deficiency leads to lanosterol and 24,25-dihydrolanosterol accumulation

To validate CYP51 depletion, we examined primary hepatocytes of iLKO and WT mice by RT-qPCR and western blot (**Figure 3**). Results showed a reduction of both *Cyp51* mRNA (**Figure 3A)** and CYP51 protein (**Figure 3B**) levels in iLKO primary hepatocytes compared to WT controls. A milder depletion of *Cyp51* mRNA was not entirely unexpected, as the cells were not cultured under cholesterol-depleted conditions that would otherwise stimulate cholesterol synthesis pathway. Instead, medium for primary cell cultivation was supplemented with 10% fetal bovine serum (FBS), which contributes to an estimated ∼30 µg/ml of cholesterol in cultivation medium [13–15].

**Figure 3.**
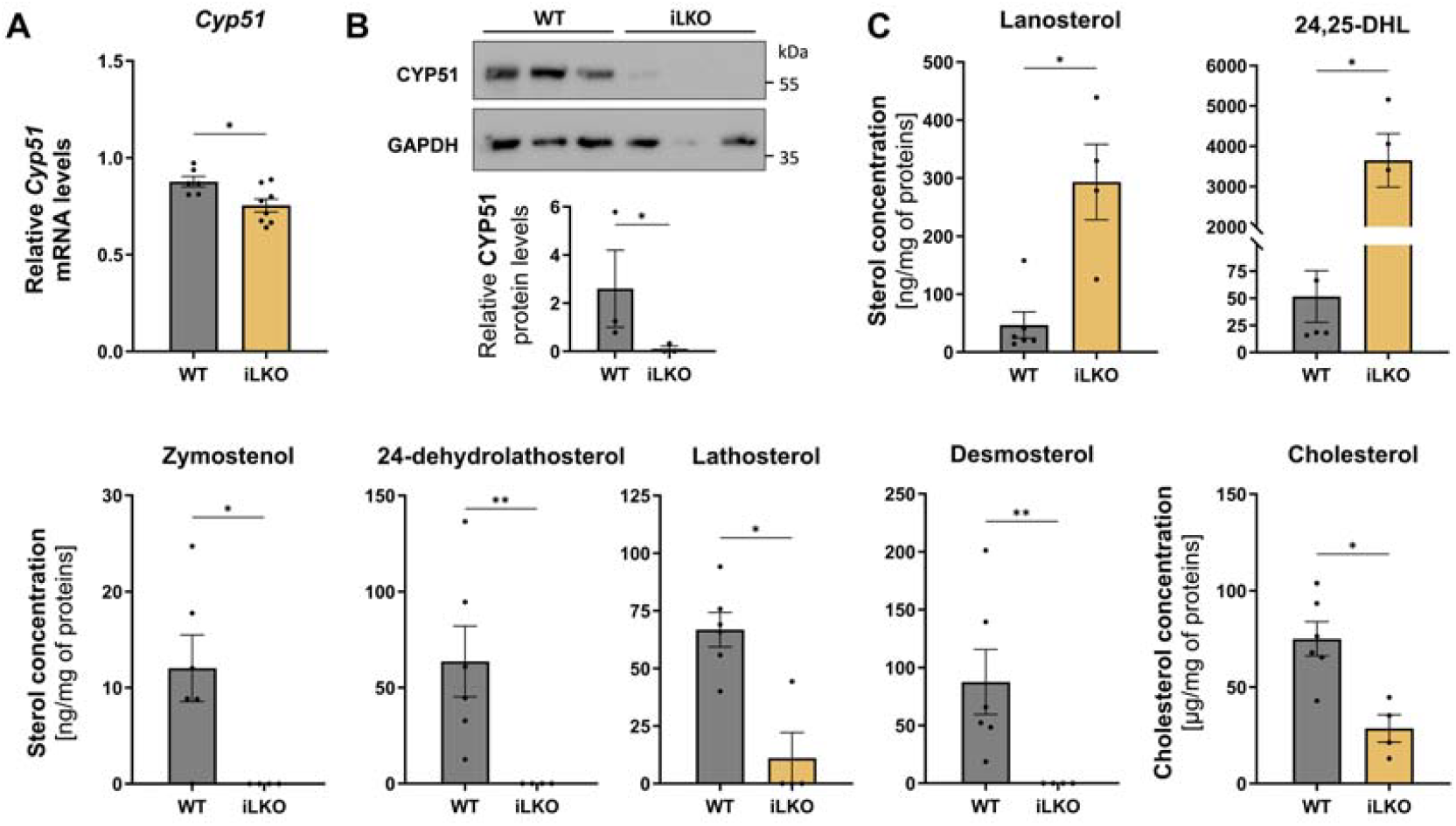
Characterization of primary hepatocytes isolated from the iLKO mouse model. **(A)** *Cyp51* mRNA expression in primary hepatocytes measured by RT-qPCR and normalized to *Utp6c, Actb* and *Hmbs* expression. Values on the graphs are means ± SEM; * p < 0.05. **(B)** Representative western blot of CYP51 in primary hepatocytes with a corresponding quantification graph. CYP51 levels were normalized to GAPDH. The mean of three iLKO and three wild type (WT) samples with SEM is presented in the plot. **(C)** Concentrations of cholesterol and sterol intermediates isolated from primary hepatocytes. All concentrations of intermediates were calculated as ng/mg or µg/mg in case of cholesterol and normalized to 100 µg of total proteins. Levels of zymosterol, 24-dehydrolathosterol, and desmosterol in iLKO samples were below detection limit. LC-MS/MS data are represented as mean ± SEM (n=4-6/group). * p < 0.05; ** p < 0.01; 24,25-DHL, 24,25-dihydrolanosterol.

We further validated *Cyp51* deletion on a functional level. As indicated in **Figure 3C**, the amount of lanosterol was 6-fold increased, while 24,25-dihydrolanosterol (24,25-DHL) was over 200-fold increased in iLKO hepatocytes compared to the WTs. Other sterols downstream of CYP51 block were either reduced (lathosterol, cholesterol) or below the detection limit (zymosterol, 24-dehydrolathosterol, and desmosterol) in iLKO primary hepatocytes (**Figure 3C**). These data confirm that the depletion of CYP51 enzyme activity was achieved after knockout induction.

Consistent with sterol profiles in isolated hepatocytes, hepatic levels of upstream CYP51 substrates were markedly elevated in iLKO livers compared to control livers (**Figure 4**). Lanosterol increased approximately 20-fold, while 24,25-DHL exhibited an even more pronounced accumulation, with more than 1000-fold increase. On the other hand, most downstream intermediates of the Bloch arm of the pathway remained unchanged, with the exception of zymosterol, which was significantly reduced, and T-MAS, which showed a slight increase in iLKO livers. Interesingly, all analysed downstream sterols of the Kandutch-Russell (K-R) pathway were significantly elevated.

**Figure 4.**
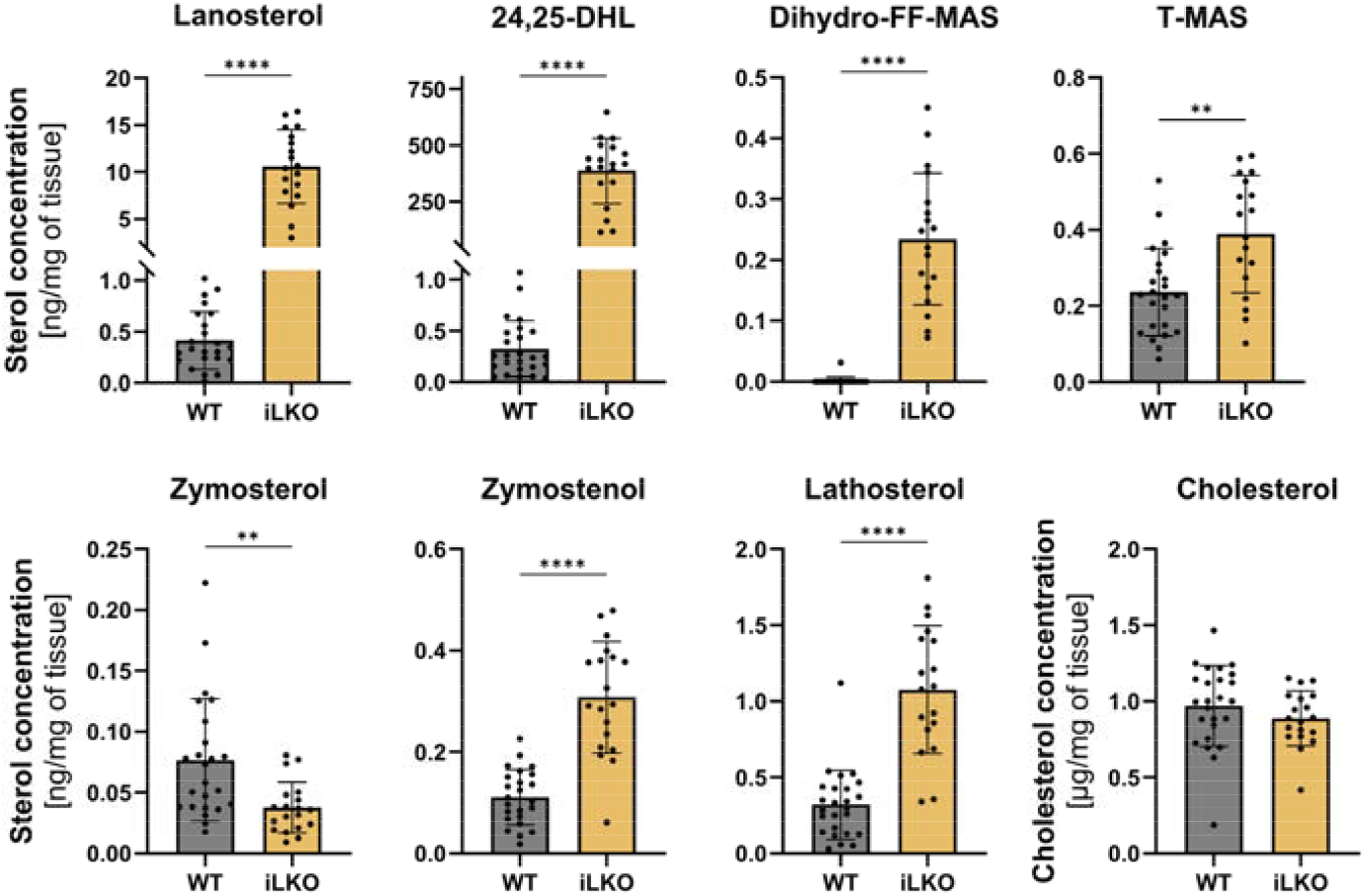
Levels of sterols extracted from liver tissue samples of iLKO and control (WT) mice. Data are pooled across both sexes and termination time points. All concentrations of intermediates were calculated as ng of sterol per mg of liver tissue or in case of cholesterol µg/mg. LC-MS/MS data are represented as mean ± SD (n=19-25/group). Mann-Whitney test was applied; only sterols with significantly changed levels (and cholesterol) are presented in this figure. ** p < 0.01; **** p < 0.0001. 24,25-DHL, 24,25-dihydrolanosterol.

To statistically evaluate the relative contributions of genotype, sex, and termination time to these sterol changes, we performed three-way ANOVA. The analysis revealed that genotype was the dominant source of variation for almost all investigated sterol levels (**Table S6, Figure S5**). For example, highly dominant was for lanosterol (76%, p < 0.0001) and 24,25-DHL (78%, p < 0.0001) and to a somewhat lesser extent for lathosterol (57%, p < 0.0001) and zymostenol (54%, p < 0.0001). In contrast, genotype explained little to no variation in downstream sterols such as desmosterol, 7-dehydrodesmosterol, and cholesterol, again indicating compensatory mechanisms. Sex × genotype interaction contributed only minor effects, which were generally small and often not statistically significant. On the other hand, time of termination exerted significant effects, although mostly minor, on several intermediates, particularly zymostenol (8%, p = 0.0001), 24-dehydrolathosterol (9%, p = 0.006), dihydro-FF-MAS (9%, p < 0.0001), and FF-MAS (18%, p= 0.003). More importantly, we observed strong genotype × time of termination interactions for multiple sterols, including zymostenol (18%, p < 0.0001), 24-dehydrolathosterol (43%, p < 0.0001), dihydro-FF-MAS (8%, p < 0.0001), zymosterol (20%, p = 0.0008) and desmosterol (31%, p < 0.0001). These sterols were upregulated in iLKO livers compared to WTs, with particularly pronounced increases in animals terminated later after *Cyp51* deletion. These findings indicate that the effects of CYP51 loss intensify over time following knockout induction, highlighting delayed metabolic rerouting within the pathway.

### 3.1. Examination of structural changes in iLKO liver

To evaluate structural changes in the liver of inducible iLKO model we performed transmission electron microscopy (TEM) in parallel with histological liver examination using hematoxylin & eosin (H&E) staining (**Figure 5, Figure S6**). Findings were compared with the previously described liver *Cyp51* Cre-loxP KO model (LKO) [7], in which the early onset of recombination presented certain limitations.

**Figure 5.**
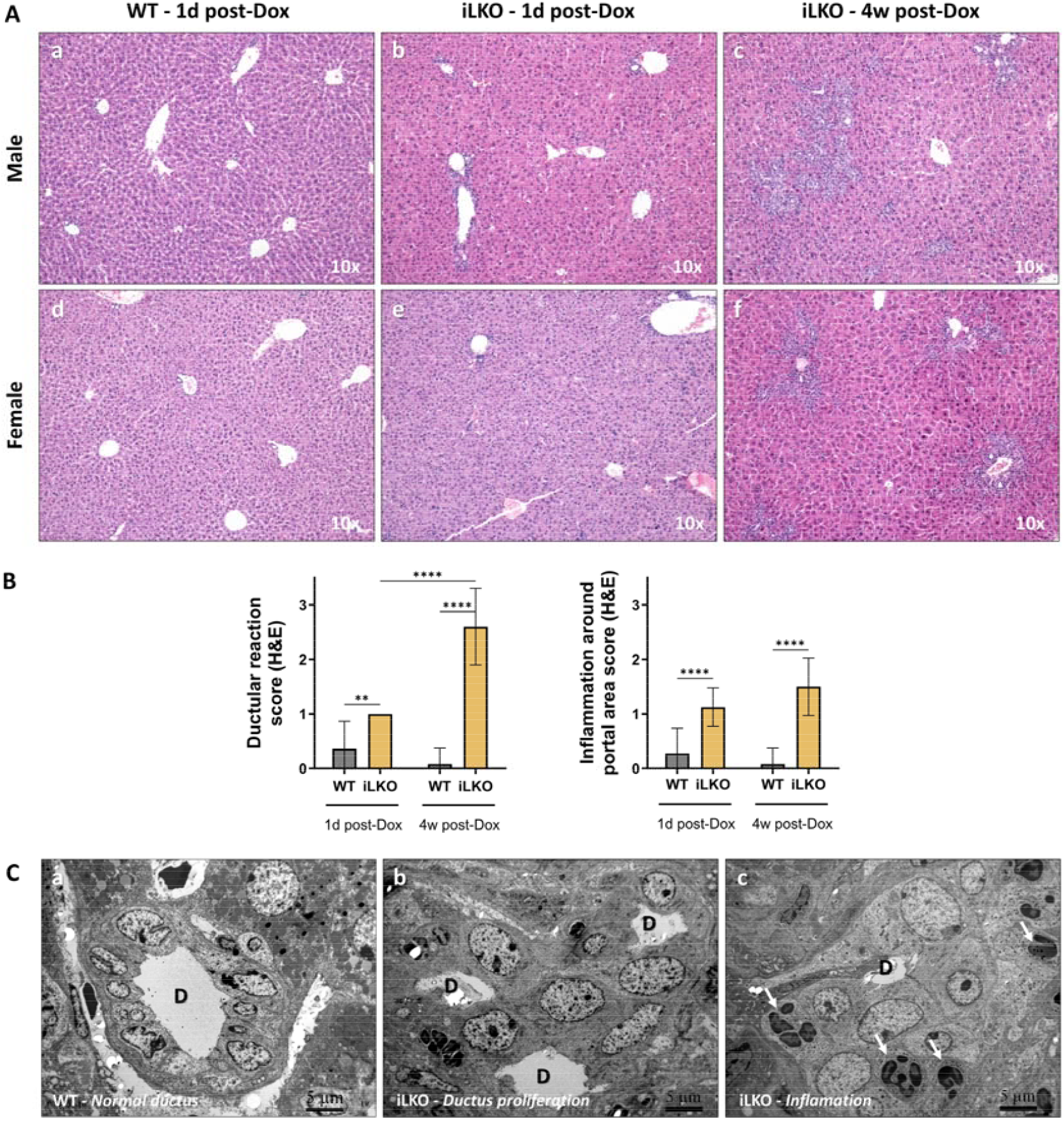
Examination of liver tissue structure. **(A)** Hematoxylin & eosin (H&E) staining representative images of normal liver in WTs (male: **a**, female: **d**), accompanied by altered livers in iLKOs, terminated 1 day (male: **c**, female: **e**) and 4 weeks (males: **d**, females: **f**) after last Doxycycline (Dox) application. 10x objective. **(B)** H&E semi-quantification data of ductular reaction and surrounding inflammation in liver samples of iLKO and WT mice, terminated 1 day or 4 weeks after last Dox application (n=10-13/group). Both processes were graded on a 0-3 scale and analyzed with two-way ANOVA and uncorrected Fisher’s test. Columns represent means ± SDs. Stat. significance is only shown for WT vs KO comparisons and between termination points.* p < 0.05; ** p < 0.01; *** p < 0.001; **** p < 0.0001. **(C)** Transmission electron microscopy (TEM) representative images: (**a**) Normal bile duct lined with cholangiocytes in WT liver tissue. (**b**) Bile duct proliferation observed in iLKO livers. (**c**) Presence of inflammatory cells (arrows) surrounding the bile duct in iLKO livers. Bile ducts are labeled as D. Figures are representative for both sexes and both termination points. Scale bar: 5 µm.

In contrast to the liver phenotype of fully developed 19-week-old LKO mice [7], the iLKO livers displayed a similar but milder phenotype, without fibrosis in either sex. Closer examination revealed the following characteristics. Normal liver structure was observed in male and female WTs **(Figure 5A: a, d)**, whereas iLKO mice exhibited mild to moderate inflammation of portal tracts, with comparable intensity at both termination time points (**Figure 5A: b, c, e, f; Figure 5B, Figure S6**). The increased oval cell proliferation (also known as the ductular reaction) was also observed mostly in iLKO livers and was more intense in the later-termination group (**Figure 5B, Figure S6**). Representative TEM images in **Figure 5C** illustrate a normal bile duct and irregular oval cell proliferation surrounded by inflammatory cells, predominantly neutrophils (arrows in **Figure 5C: c**). Interestingly, a three-way ANOVA of H&E and TEM data indicated that sex was not a significant main effect for neither inflammation nor ductular reaction (data not shown), suggesting that these processes are less likely involved in female-pronounced hepatomegaly (**Figure 2B**).

To further investigate the mechanisms underlying these changes, we examined subcellular structures by TEM. While WT livers showed mitochondria that were mostly intact to slightly swollen and endoplasmic reticulum (ER) with preserved morphology (**Figure 6A**), iLKO livers exhibited more pronounced mitochondrial swelling (**Figure 6B**) and dilated ER cisternae (**Figure 6C**). In addition, multilamellar structures/myelin-like figures were detected in some iLKO hepatocytes (**Figure 6D**). No significant differences were observed between male and female or between different termination points. However, visual assessment, while unbiased (blinded), would benefit from confirmation by quantitative image analysis. Overall, iLKO livers displayed only slight alterations in organelle structures when compared with WT.

**Figure 6.**
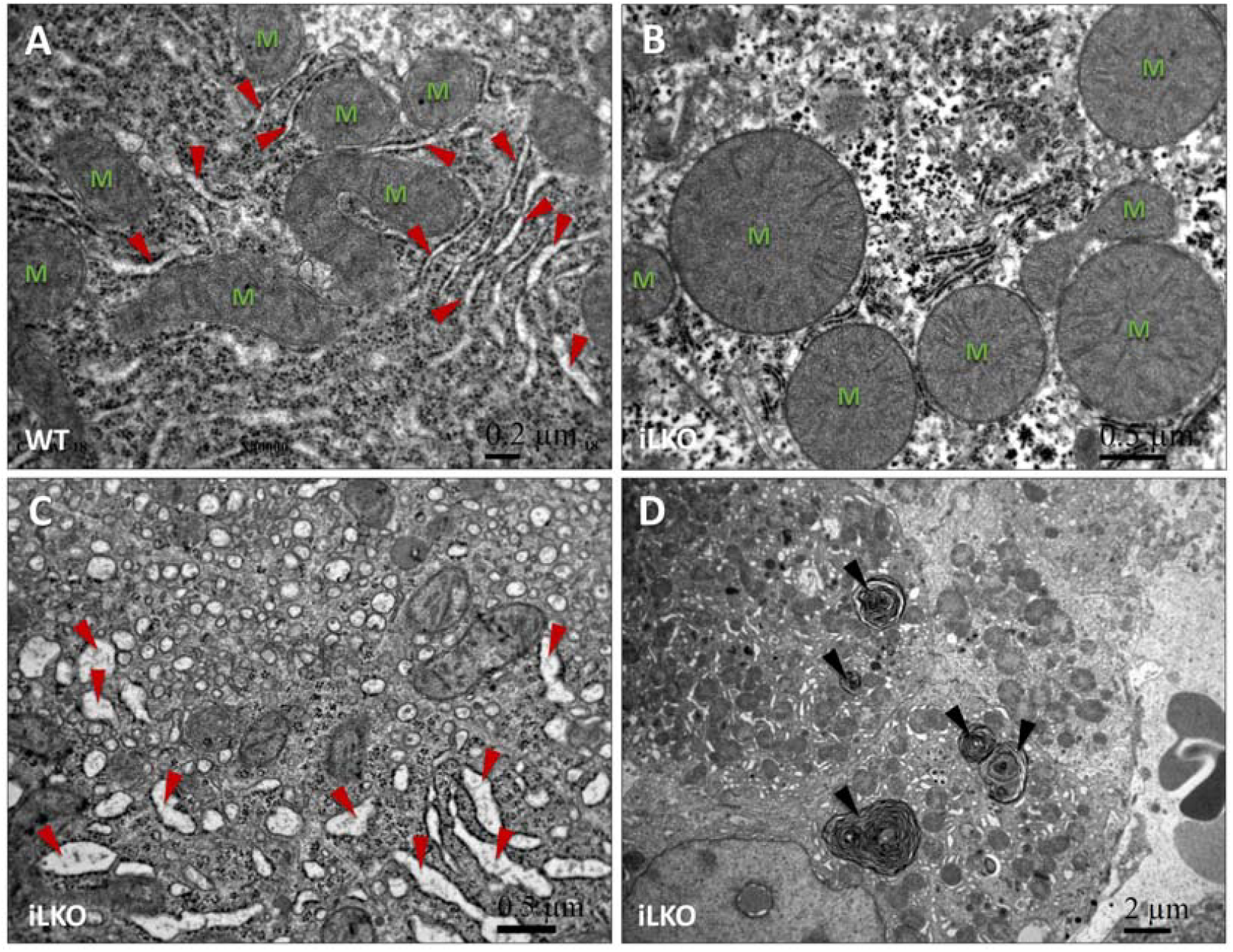
Transmission electron microscopy (TEM) of iLKO and wild type (WT) livers. **(A)** Mitochondria and endoplasmic reticulum (ER) of WT hepatocytes. **(B)** Slightly swollen mitochondria in an iLKO hepatocyte. **(C)** Dilated ER in an iLKO hepatocyte. **(D)** Multilamellar structures in iLKO hepatocytes. Labels: N – nucleus (blue); M – mitochondrion (green); red arrowheads – ER, black arrowheads - multilamellar structures. Scale bars: 0.2 µm (A), 0.5 µm (B, C), 2 µm (D). Figures are representative of both sexes and both termination times.

### 3.4. Disturbed sterol metabolism promotes nuclear lipid droplet formation

Beyond the subtle structural changes observed in iLKO hepatocytes, a more pronounced alteration was the presence of lipid droplets (LDs). LDs are essential organelles that store neutral lipids and help maintain cellular lipid and energy homeostasis. Originating from the ER, they dynamically interact with other organelles and buffer toxic lipid species, making them central hubs of metabolic regulation [16]. Since cholesterol synthesis is tightly linked to lipid metabolism, we investigated whether *Cyp51* deletion affects hepatic LD accumulation.

Examination of liver sections by H&E straining revealed the presence of LDs in both WT and iLKO hepatocytes, but with no significant differences in their quantity between sample groups (**Figure S7**). The TEM analysis similarly showed no significant changes in total LD numbers, however, iLKO liver samples exhibited a higher abundance of nuclear LDs in comparison to controls (**Figure 7B; Figure 7C: a, b)**. This difference was statistically significant in 4 weeks post-Dox group, showing not only higher LD presence compared to WT, but also compared to 1 day post-Dox iLKO samples. This finding suggests that nuclear accumulation of LDs is a delayed response to *Cyp51* deletion. Three-way ANOVA indicated that sex was not a significant main effect (data not shown). Closer examination showed varying degrees of heterochromatin proximity to LDs (**Figure 7C: c**), similar as already observed in human liver samples [17]. These findings support the view that LD-chromatin interactions are a conserved feature potentially contributing to nuclear organization and gene regulation, and it would be of interest to further evaluate whether such proximity differs between WT and iLKO samples using advanced image analysis tools [18–20].

**Figure 7.**
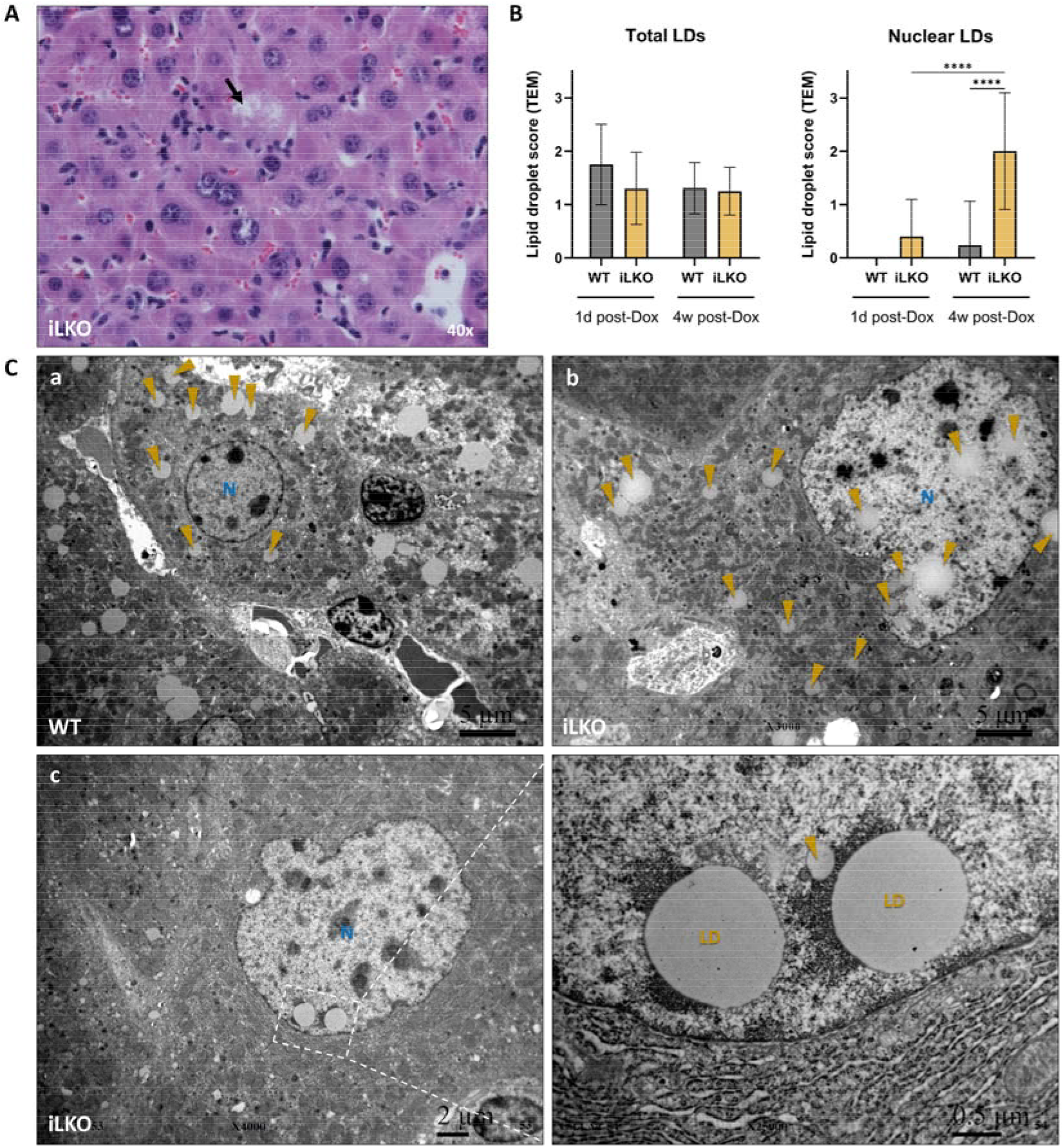
Presence of lipid droplets (LDs) in iLKO and WT liver. **(A)** Hematoxylin & Eosin (H&E) staining of iLKO liver with visible lipid droplets (black arrow) in hepatocytes; 40x objective. **(B)** Transmission electron microscopy (TEM) semi-quantification data of LD presence graded on a 0-3 scale and analyzed with ordinary two-way ANOVA and uncorrected Fisher’s test. Samples were grouped based on genotype (WT or iLKO), and time of termination (1 day or 4 weeks post doxycycline (Dox) treatment). Columns represent means ± SDs (n=10-16/group); **** p < 0.0001. **(C)** Representative TEM images: **(a)** LDs in a hepatocyte of WT liver, bar: 5 µm. **(b, c)** iLKO hepatocyte with cytoplasmic and nuclear LDs; bars: 5 µm (**b**), 2 µm and 0.5 µm (**c**). Representative for both sexes and both termination points. Labels: N (blue) – nucleus; LD or arrowheads (yellow) - lipid droplets.

### 3.5. Accumulation of sterol intermediates may lead to crystal formation in CYP51-deficient liver cells

Surprisingly, with the TEM we discovered a formation of crystal-like structures and acicular clefts (elongated, needle-like spaces) exclusively in iLKO liver samples of both sexes and ages (**Figure 8A: a, b**). No similar structures were found in the WTs. A detailed examination showed abundant rectangular and oval-shaped crystals especially within enlarged Kupffer cells of iLKO livers (**Figure 8A: c**). The crystals varied in size, with median width 89 nm (95% CI: 66-133) and median length 918 nm (95% CI: 573-1200) (**Table S7**). Semi-quantitative TEM analysis (**Figure 8B**) showed significantly increased presence of crystal structures in hepatocytes and Kupffer cells of iLKO livers. Three-way ANOVA indicated no significant main effect of sex (data not shown), so samples were pooled accordingly. In contrast, time of termination had a significant main effect, suggesting that crystal formation represents a delayed response to *Cyp51* deletion. Finding of these structures may be also related to the increased number of apoptotic cells observed in the liver parenchyma of iLKO mice (**Figure 8C**).

**Figure 8.**
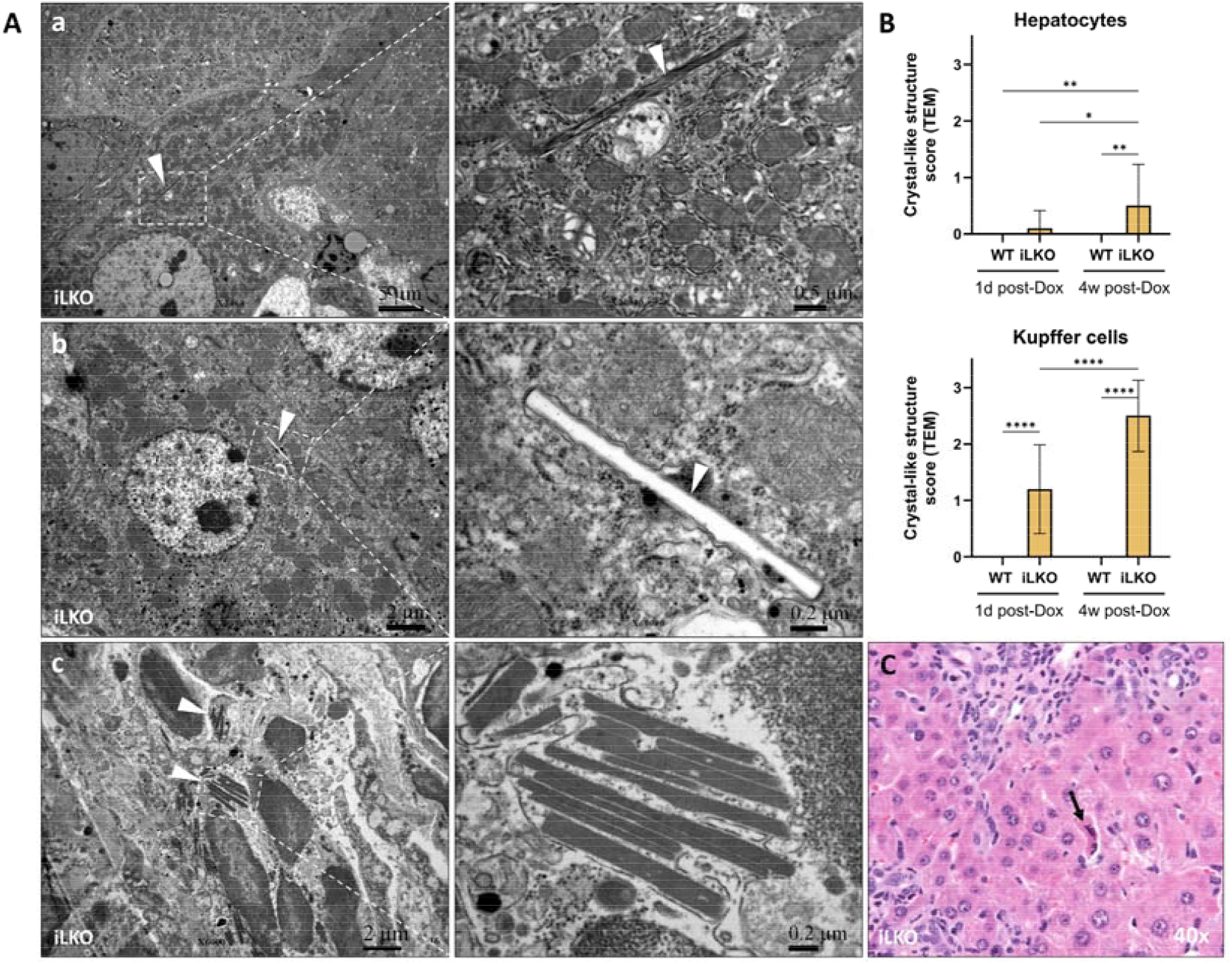
**(A)** Transmission electron microscopy (TEM) of iLKO livers revealed presence of crystal-like structures of unknown composition. **(a)** Example of a crystal (bars: 5 µm and 0.5 µm) and **(b)** an acicular cleft (bars: 2 µm and 0.2 µm) in hepatocytes, both indicated with white arrowheads. **(c)** Rectangular crystals in the cytoplasm of Kupffer cells; bar: 2 µm and 0.2 µm. **(B)** Quantified TEM data of crystals in the liver of iLKO and WT mice, 1 day or 4 weeks post doxycycline (Dox) treatment (n=10-16/group). Presence of structures was graded on a 0-3 scale and analyzed with ordinary two-way ANOVA and uncorrected Fisher’s test. Columns represent mean ± SD. * p < 0.05; ** p < 0.01; **** p < 0.0001. **(C)** Example of an apoptotic iLKO hepatocyte (black arrow), which were widely distributed in the liver parenchyma. Objective 40x.

To characterize the composition of the crystals identified in the livers of iLKO mice, we performed matrix-assisted laser desorption ionization–time of flight mass spectrometry imaging (MALDI-TOF MSI) analysis. Based on the LC-MS/MS results, which showed a high accumulation of lanosterol and 24,25-DHL in the iLKO livers (**Figure 3C and 4**), we hypothesized that the crystals may be composed of these two sterols. However, the crystals were too small to be reliably localized using light microscopy on liver tissue sections as well as signal spatial resolution of MSI was not sufficient to be able to show correlation of signal and crystal overlay. An additional obstacle were possible isobaric sterols that share the same monoisotopic mass but differ structurally. **Figure S8** shows homogeneously dispersed 24,25-DHL in one iLKO and one WT liver section. Although the intensity is much higher in the iLKO sample, the technical limitations of our MALDI-TOF MSI system made the determination of the crystal composition unfeasible.

Interestingly, the formation of crystals appears to be a specific feature of iLKO hepatocytes *in vivo*, as similar structures were not observed in a liver cell line. Despite comparable accumulation of lanosterol and 24,25-DHL in human hepatoblastoma HepG2 cells with depleted CYP51 [10], no crystals were detected by TEM (**Figure 9**). Instead, these cells exhibited a noticeable increase in LDs compared to control HepG2 cells, with LDs present in both the cytoplasm and nucleus (**Figure 9F**). No visible alterations were observed in the structure of the ER or mitochondria (**Figure 9C–E**).

**Figure 9.**
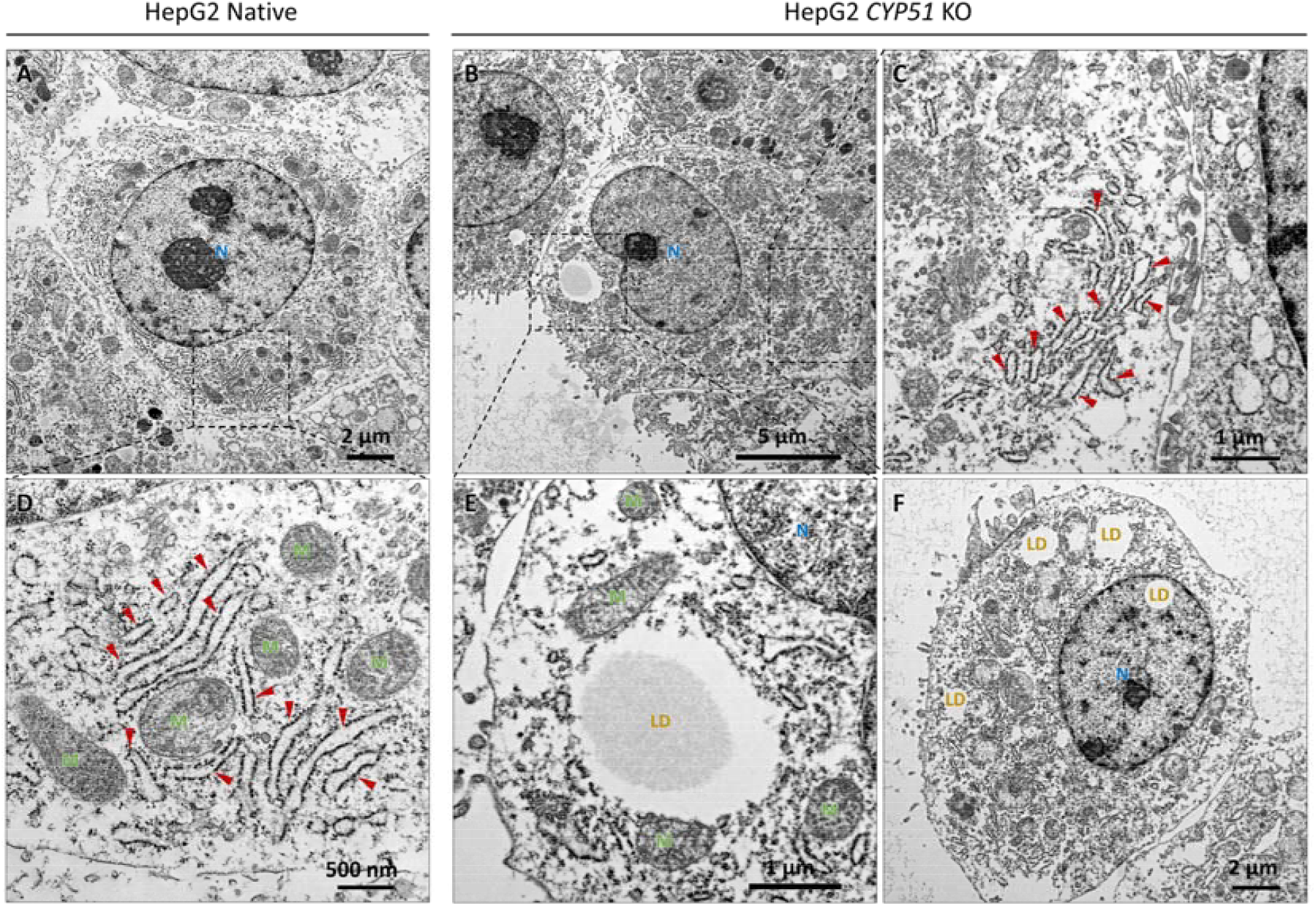
Transmission electron microscopy (TEM) representative images of HepG2 *CYP51* KO cell model (right) and its control HepG2 Native cell line (left). **(A, D)** HepG2 Native cell with magnified region showing endoplasmic reticulum (ER) and mitochondria. **(B, C, E)** HepG2 *CYP51* KO cell with magnified region showing ER, mitochondria and a lipid droplet (LD). **(F)** HepG2 *CYP51* KO cell with LDs present both in cytoplasm and nucleus. Labels: N – nucleus (blue); M – mitochondrion (green); LD – lipid droplet (yellow); red arrowheads – endoplasmic reticulum. Scale bars: 5 µm (B), 2 µm (A, F), 1 µm (C), 500 nm (D).

## 4. Discussion

Here we introduce an inducible, liver-specific *Cyp51* knockout mouse model (iLKO) that circumvents developmental confounders of albumin-Cre model LKO [7] and, yields adult primary hepatocytes appropriate for further *in vitro* investigation. By inducing recombination in mature liver, iLKO separates consequences of acute sterol disequilibrium from those due to altered liver development. Although, iLKO mice exhibited largely similar hepatic phenotype as adult LKO model mice [7], with developed hepatomegaly, oval cell proliferation and inflammation, fibrosis was not as apparent. Therefore, otherwise unsuccessful primary hepatocyte isolation was now possible.

As expected from blocking CYP51, iLKO primary hepatocytes and liver tissue accumulated upstream substrates, lanosterol and even more strongly DHL. The latter, showed striking ∼200-fold increase in hepatocytes and ∼1000-fold in hepatic tissue. Similar findings were seen in other *Cyp51* KO models [4,5,7,8,10]. The greater accumulation of DHL relative to lanosterol, is consistent with the expected more prominent enzymatic conversion by 24-dehydrocholesterol reductase (DHCR24) under conditions of CYP51 inhibition. Concomitantly, downstream sterols were reduced in primary hepatocytes, while tissue analysis indicated flux rerouting toward the K-R arm (elevated K-R intermediates, reduced zymosterol and unchanged Bloch pathway intermediates). A likely interpretation is that the hepatic cells in which synthesis remains active take up accumulated DHL and then cholesterol synthesis proceeds via the K–R pathway. Total hepatic cholesterol remained unchanged, implying compensation (e.g., uptake from other cells/tissues) that buffers cholesterol levels despite the biosynthesis block in hepatocytes.

At the tissue level, iLKO showed mild portal inflammation and a ductular reaction without overt fibrosis at the studied time points (age and/or time of examination after KO induction). This supports an earlier stage of hepatic injury, observed in previous *Cyp51* KO models [7,8]. Notably, sex was not a significant driver of inflammation or ductular reaction in this setting. Subcellular hepatocyte structures were mostly only slightly altered in iLKO mice, which showed a more pronounced mitochondrial swelling and dilated ER compared to WT mice. The latter is consistent feature of ER stress [21], and resembles findings from previous *Cyp51* KO or inhibition models that reported upregulation of unfolded protein response genes (*Casp12, Eif2ak3, Eif2s1, Eif2ak4*) and associated transcription factors (NFY) [4,7,22]. Since the KO in those mouse models occurred before liver maturation, our findings indicate that the previously observed ER stress reflects a genuine consequence of CYP51 loss rather than a developmental defect caused by early deletion. Interestingly, HMG-CoA reductase (HMGCR) KO models also display ER dilation, along with ER stress-induced apoptosis [23,24]. Similarly, HMGCR is expected to be reduced in our model [7] via Insig-dependent ER-associated protein degradation (ERAD) as a result of lanosterol and DHL accumulation [25–27]. Since lipotoxicity is already recognized inducer of hepatic ER stress [28,29] we hypothesize that, in our model, it manifests as an accumulation of lanosterol and DHL, which might trigger ER stress by altering HMGCR levels or, more broadly, by interfering protein synthesis and processing in ER. The latter could occur similarly as the effects of ER-cholesterol build-up [30,31], however, because cholesterol and its precursors differentially affect membrane properties [32–35], this calls for further mechanistic investigation.

In addition to subtle alterations observed in organelle structures, more pronounced was accumulation of LDs in nuclei of iLKO hepatocytes, without a change in total LD counts. Nuclear LDs were first observed in hepatocytes more than 60 years ago [36], but their origin has only more recently begun to be understood [37,38]. Three forms of LDs have been recorded in hepatocytes nuclei and are generally distinguished by whether they are surrounded by inner and outer nuclear membranes, only the inner membrane, or lack a membrane entirely [17,37,39]. First, cytoplasmic LDs, are not a true nuclear LD type, but instead result from cytoplasmic invaginations into the nucleus, therefore can be distinguished by being enclosed by both nuclear membranes (i.e. within type II nucleoplasmic reticulum (NR)). Second group are formed in the ER lumen, under certain conditions enlarge and accumulate in type I NR that can release them after its breakage. These are therefore referred to as NR-lumenal LDs. The third, nucleoplasmic LDs, lack surrounding nuclear membranes and are thought to arise from NR-lumenal LDs, after membrane disintegration. More detailed investigation proposed that NR-lumenal LDs accumulate under ER stress and once they become nucleoplasmic, these LDs recruit CTP:phosphocholine cytidylyltransferase α (CCTα), the rate-limiting enzyme of phosphatidylcholine (PC) synthesis [39]. This stimulates *de novo* PC production, which has been proposed as a feedback mechanism to alleviate ER stress through ER expansion [40]. In our study, nuclear LDs were markedly increased in CYP51-deficient livers, and based on the above work we hypothesize that their accumulation reflects ER stress triggered by sterol imbalance. The predominance of the nucleoplasmic form in our samples, suggested by the TEM detection of LDs lacking membranes, further supports this idea. Similar associations have been observed clinically. Imai et al. [41] reported that nucleoplasmic LDs are a common feature in liver biopsies from patients with autoimmune hepatitis (AIH), drug-induced liver injury (DILI), and metabolic dysfunction-associated steatohepatitis (MASH; formerly known as nonalcoholic steatohepatitis, NASH). In the same study, hepatocytes with minimal histological changes rarely contained nuclear LDs, whereas their frequency increased in livers showing enlargement and evidence of ER stress. Additionally, in our study, nuclear LD accumulation became significant only in a 4 weeks post-Dox group, indicating that nuclear LDs may be a delayed adaptation, associated with increased hepatomegaly. Despite these mechanistic links, hepatic nuclear LDs could possibly arise by a different mechanism as demonstrated in other mammalian cell types [37].

The most surprising observation in our study is the presence of crystal-like structures (in this manuscript referred to as crystals), exclusively in iLKO liver tissue. Similar inclusions have been described in livers of patients with cholesteryl ester storage disease (CESD) [42] and MASH [28,43–46], where they are composed of free cholesterol. Cholesterol crystals have been studied extensively in atherosclerosis [47–56], where they have been shown to promote lesion development by activation of the potent Nod-like receptor protein (NLRP)3 inflammasome [50,57]. Similar findings were reported for inflammation occurring in the uterine wall [58] as well as in MASH [28,43,44]. The precise mechanism of hepatic cholesterol crystal formation and connection to MASH remains unresolved, however Ioannou et al. [43–46] proposed a model based on their work. They suggest that cholesterol accumulates within hepatocyte LDs, where excess free cholesterol crystallizes out of the LD’s phospholipid monolayer when reaching a saturation threshold. Further, necrotic hepatocytes become encircled by Kupffer cells that phagocytose and enzymatically degrade remnant LDs. Exposure to cholesterol crystals causes activation of the NLRP3 inflammasome within Kupffer cells, which leads to production of proinflammatory cytokines and chemokine, sustaining sterile inflammation typical of MASH. Finally, these chemotactic signals attract an inflammatory infiltrate of additional Kupffer cells and neutrophils, as well as causing aggregation, activation, and transformation of stellate cells into collagen-producing myofibroblasts, leading to fibrosing MASH. Additionally, Bashiri et al. [28] also opened a question of possible connection of cholesterol crystal with ER stress.

In our iLKO model, the crystals are unlikely to be cholesterol-based, as neither hepatocyte nor hepatic tissue cholesterol was elevated. Instead, given the marked accumulation of lanosterol and DHL, we hypothesize that the crystals comprise lanosterol and/or DHL. To our knowledge, non-cholesterol sterol crystals have not yet been reported in mammalian tissues. Crystals also were not identified in LKO models [4,7,8], as those studies did not use TEM or other high-resolution imaging. Although we were unable to confirm crystal composition due to technical limitations, several observations support their pro-inflammatory role: they were predominantly localized within Kupffer cells rather than hepatocytes, increased oval-cell proliferation was observed with portal tracts exhibiting neutrophil-rich inflammation. Moreover, crystalline particulates of endogenous or exogenous origin are well known to trigger inflammation [59]. To determine whether crystallization is a general consequence of precursor-sterol overload, we evaluated HepG2 *CYP51* KO cells. Despite the accumulation of lanosterol and DHL [10], no crystals were detected by TEM. This negative result may reflect the cancer-derived nature of HepG2 cells (altered metabolic and cellular processes), the absence of native liver architecture (3D organization, non-parenchymal cell types, extracellular matrix etc.), or simply culture conditions that did not reach the crystallization threshold. In contrast, Ioannou et al. [45] reported that exposure of HepG2 cells to low-density lipoprotein (LDL)-cholesterol plus oleic acid induced LD-associated cholesterol crystals that activated co-cultured THP-1 macrophages. Notably, the crystals only appeared after 20 days of treatment, similarly as the crystals observed in iLKO liver were more frequent in the 4 weeks post-Dox group. This suggests that non-sterol crystals might also occur in HepG2 *CYP51* KO cells when cultured under sterol-accumulating conditions for a prolonged period. Together, these findings raise the possibility that sterol crystal–driven innate immune activation extends beyond cholesterol *per se* and may contribute to disease progression in the *Cyp51* KO models [8].

The observed accumulation of CYP51 substrates in iLKO livers prompts the question of their fate. As they are known to be cytotoxic [35], cells must find ways to dispose of them. The accumulated nuclear LDs could suggest that these serve for compartmentalization of toxic sterol intermediates. However, lanosterol and DHL are poor substrates for acyl CoA:cholesterol acyltransferase (ACAT) [60,61] and thus are unlikely to be efficiently esterified and sequestered in LD cores; at most, they may transiently partition into their phospholipid monolayer. Other possible route is their secretion into bile. Analyses of human gallbladder bile and gallstones have identified lanosterol (and to lesser extend DHL) among their sterol constituents and shown that methylsterols sediment upon bile centrifugation [62]. Consistent with this, the sand-like material observed in the bile of prematurely euthanized iLKO subjects of our study likely reflects elimination of excess lanosterol via secretion into the bile. This interpretation again parallels cholesterol crystallization, as cholesterol crystals are found in supersaturated bile of patients with cholelithiasis and are linked to gallstone formation and gallbladder inflammation [63]. Another way to eliminate excess lanosterol is ATP-binding cassette transporter A1 (ABCA1)-dependent efflux from the plasma membrane to extracellular apoA-I, forming nascent high-density lipoproteins (HDL). ABCA1 preferentially releases newly synthesized C29/C30 precursor sterols (including lanosterol) over cholesterol, providing a possible rapid detoxification route [64]. The crystals observed in iLKO Kupffer cells further suggest that these cells may acquire sterol intermediates from hepatocytes. Beyond phagocytosis, ABCA1-dependent lanosterol efflux coupled with HDL uptake by Kupffer cells [65] offers a plausible transfer pathway. Although it is unclear how excess sterol amounts affect Kupffer cells, Buttke et al. [66] showed that effects of methylsterols can even be favorable in some cell types.

### 4.1. Study limitations and future directions

Despite providing valuable insights into CYP51 depletion and sterol buildup in the liver, our study also holds some limitations. First, we did not include a control group without Dox application in order to keep the animal/sample numbers manageable. However, prior work using the same parental mouse line, LC-1/rTA^LAP^-1, and Dox application protocol has already shown that Dox does not affect body weight or the expression of key lipid metabolism genes in hepatocytes [67,68]. The crystal-like inclusions seen here warrant chemical identification, which was not possible with our current tools. In addition, sterol and crystal accumulation may elicit sex-specific responses. It would also be useful to examine how CYP51-deficient hepatocytes interact with other liver cells, because sterol profiles differed between whole tissue and isolated hepatocytes, and crystal inclusions were mainly present in Kupffer cells. Similarly, there is also a need to characterize nuclear LDs by assessing their lipid composition [69,70], LD-associated proteins [71,72], markers of ER stress [21,73], interactions of LDs with other organelles and nuclear domains [69,74,75] etc. The link between sterol intermediates and the observed hepatic changes could be further verified with additional interventions, such as high dietary cholesterol [7], lanosterol supplementation, application of statins or small-molecule CYP51 inhibitors [22]. Notably, our semi-quantitative TEM scoring might have missed some organelle changes. More objective approaches would therefore provide more reliable effect observations. With non-hepatic tissues already collected, it will also be possible to map systemic changes in precursor sterols, organelle stress, and inflammation across organs. Finally, transcriptomic and proteomic analysis of the iLKO model could map pathway changes beyond cholesterol synthesis, including bile acid and steroid pathways. This would also allow for better comparison with human data and would help place our findings in a broader disease context (e.g. metabolic syndrome, MASH, cancer).

## 5. Conclusions

We established an inducible, liver-specific *Cyp51* knockout mouse model (iLKO) that enables adult-onset deletion and useful isolation of primary hepatocytes. iLKO exhibits lanosterol and 24,25-DHL accumulation and unchanged or mildly reduced cholesterol levels. Livers show mild portal inflammation, ductular reaction, ER dilation, and mitochondrial swelling, and delayed accumulation of nuclear LDs. We also observed crystal-like inclusions, especially in Kupffer cells, suggesting a novel possible mechanism triggering inflammation due to excess toxic precursors. iLKO thus provides a practical platform to dissect how specific sterol intermediates drive organelle stress and innate immune activation; future work should define crystal composition, inflammasome engagement, and sterol flux into other metabolic pathways.

## Acknowledgements

The authors thank prof. dr. Rolf Gebhardt from the Institute of Biochemistry, Faculty of Medicine, University of Leipzig, Germany, for rTA_LAP_-1/LC1-1 transgenic mice gifted to prof. dr. Damjana Rozman for experimental research of Cyp51 liver function, and to Petra Nassib from the Centre for Functional Genomics and Bio-Chips, Institute of Biochemistry and Molecular Genetics, Faculty of Medicine, University of Ljubljana, for technical support. We would also like to acknowledge the use of <u>BioRender</u> for creation of graphical abstract and indicated figures, and GraphPad Prism (version 10.3.1 for Windows) for statistical analysis and data visualization.

## CRediT authorship contribution statement

Tinkara Kreft: Methodology, Investigation, Formal analysis, Validation, Visualization, Writing – original draft, Writing – review & editing. **Kaja Blagotinšek Cokan**: Conceptualization, Methodology, Investigation, Formal analysis, Visualization, Writing – original draft, Writing – review & editing. **Cene Skubic**.: Methodology, Investigation, Formal analysis. **Tadeja Režen**: Resources, Writing – review & editing. **Martina Perše**: Methodology, Investigation, Supervision. **Karmen Wechtersbach**: Investigation, Writing – review & editing. **Nika Kojc**: Investigation. **Jera Jeruc**: Investigation, Supervision, Writing – review & editing. **Željko Debeljak**: Investigation, Formal analysis. **Marija Heffer:** Investigation, Supervision. **Madlen Matz-Soja:** Methodology, Supervision, Resources. **Damjana Rozman**: Conceptualization, Supervision, Resources, Project administration, Funding acquisition, Writing – original draft, Writing – review & editing.

## Funding

This research was funded by the Slovenian Research and Innovation Agency (ARIS) grants P1-0390, J1-9176, J1-50024, IP-0022 (CFGBC), ESFRI-ELIXIR, BI-DE/17-19-8, and the ARIS Young Researcher grants for K. Blagotinšek Cokan and T. Kreft. Study was also funded by the Federal Ministry of Education and Research (BMBF, Germany) within the research network LiSyM [031L0053] and LiSyM Cancer [031L0256C and 031L0258E].

## Conflicts of Interest

None.

## Declaration of generative AI and AI-assisted technologies in the writing process

During the preparation of this work, the authors used ChatGPT (OpenAI; GPT-5) in order to improve the readability and language of the manuscript. After using this tool, the authors reviewed and edited the content as needed and take full responsibility for the content of the published article.

## 7. Supplementary material

## Notes

### Competing Interest Statement

The authors have declared no competing interest.

