## Supplementary material for "Hepatic management of toxic sterols after acute deletion of *Cyp51* from cholesterol synthesis"

Research article

**Supplementary materials:**

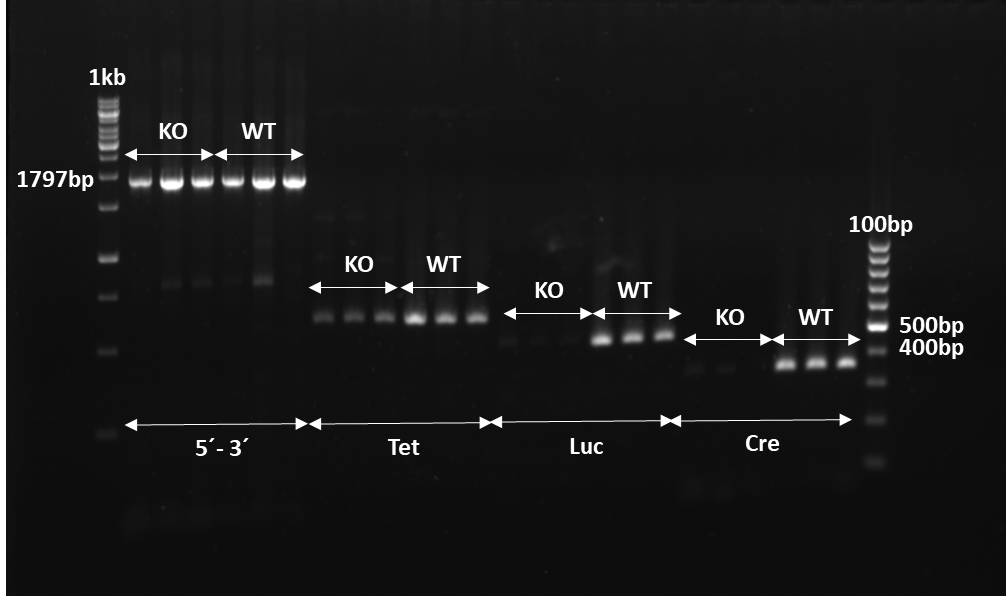

**Figure S1**. Ear clip gDNA genotyping with 5lox-3lox, Tet, Cre, and Luc primers. iLKO (i.e. KO) genotype is confirmed by the presence of all 4 analyzed fragments: 5lox - 3lox (1797 bp), Tet (500 bp), Luc (500 bp) and Cre (400 bp). The presence of 5´-3´ and tet fragments and absence of Cre and Luc fragments indicate WT genotype.

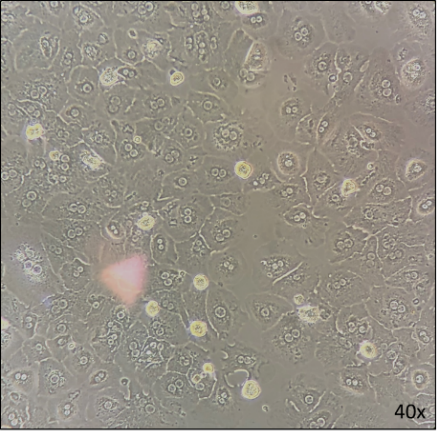

**Figure S2.** Primary hepatocyte plating – after 24 h acquired their typical hexagonal shape as seen under light microscopy (Olympus Microscope IX2-SLP); 40x objective.

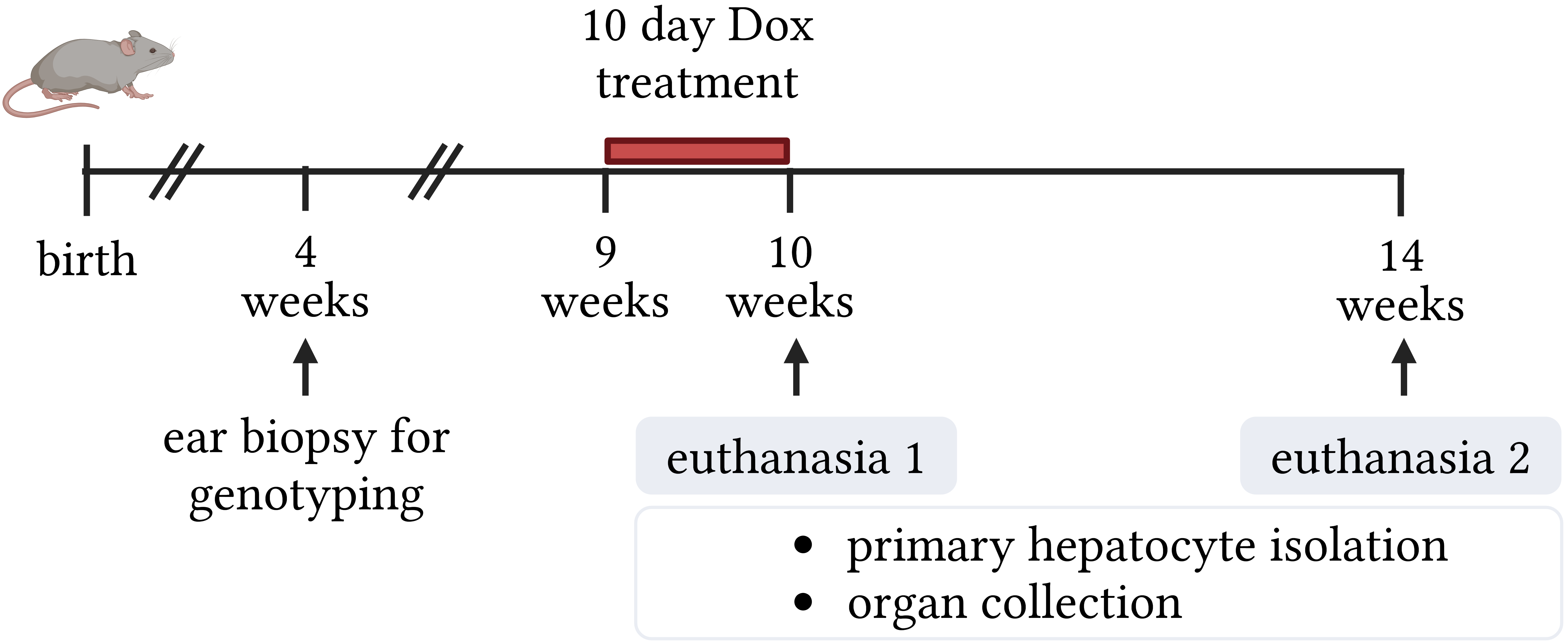

**Figure S1.** Experimental design. At 4 weeks of age ear biopsies were taken for genotyping by PCR. Both iLKO mice and wild-type (WT) mice were included in the subsequent procedures. At 9 weeks of age, mice received doxycycline (Dox; 2 mg/ml) in drinking water for 10 days to induce recombination with the CRE-loxP system. One half of the subjects was euthanized one day after the last Dox application (at ~10 weeks of age). The second half was euthanized 4 weeks later (at ~14 weeks of age), to evaluate delayed effects. The WT mice served as a control to distinguish the effects of Cyp51 deletion from potential off-target effects of doxycycline treatment or the genetic background. At termination, a subset of mice was used for primary hepatocyte isolation, and the remainder for organ collection, with the present manuscript focusing on analysis of the liver. Figure was created with BioRender.

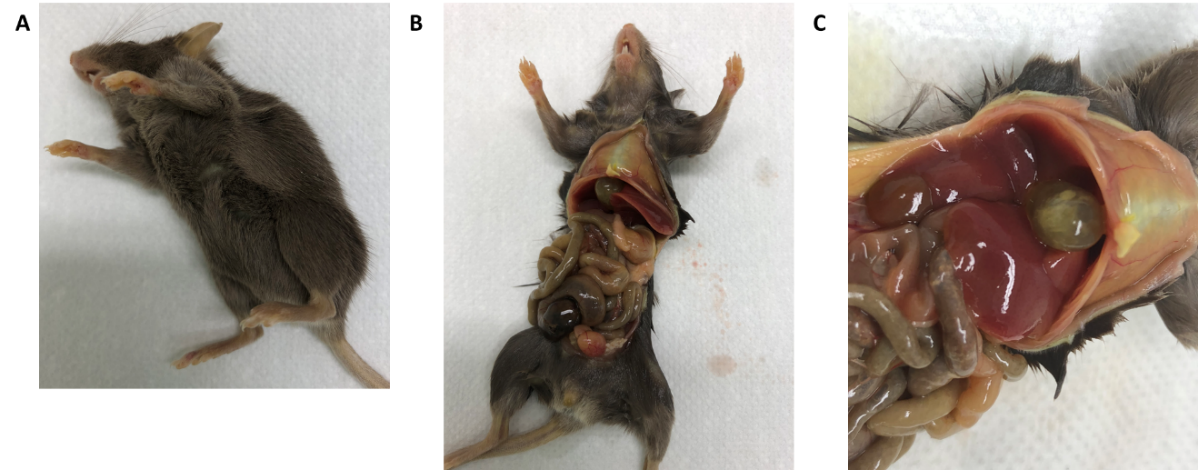

**Figure S2 .** iLKO mice euthanized between 11-13 weeks of age due to changed posture, apathetic behaviour and the suspicion to jaundice.

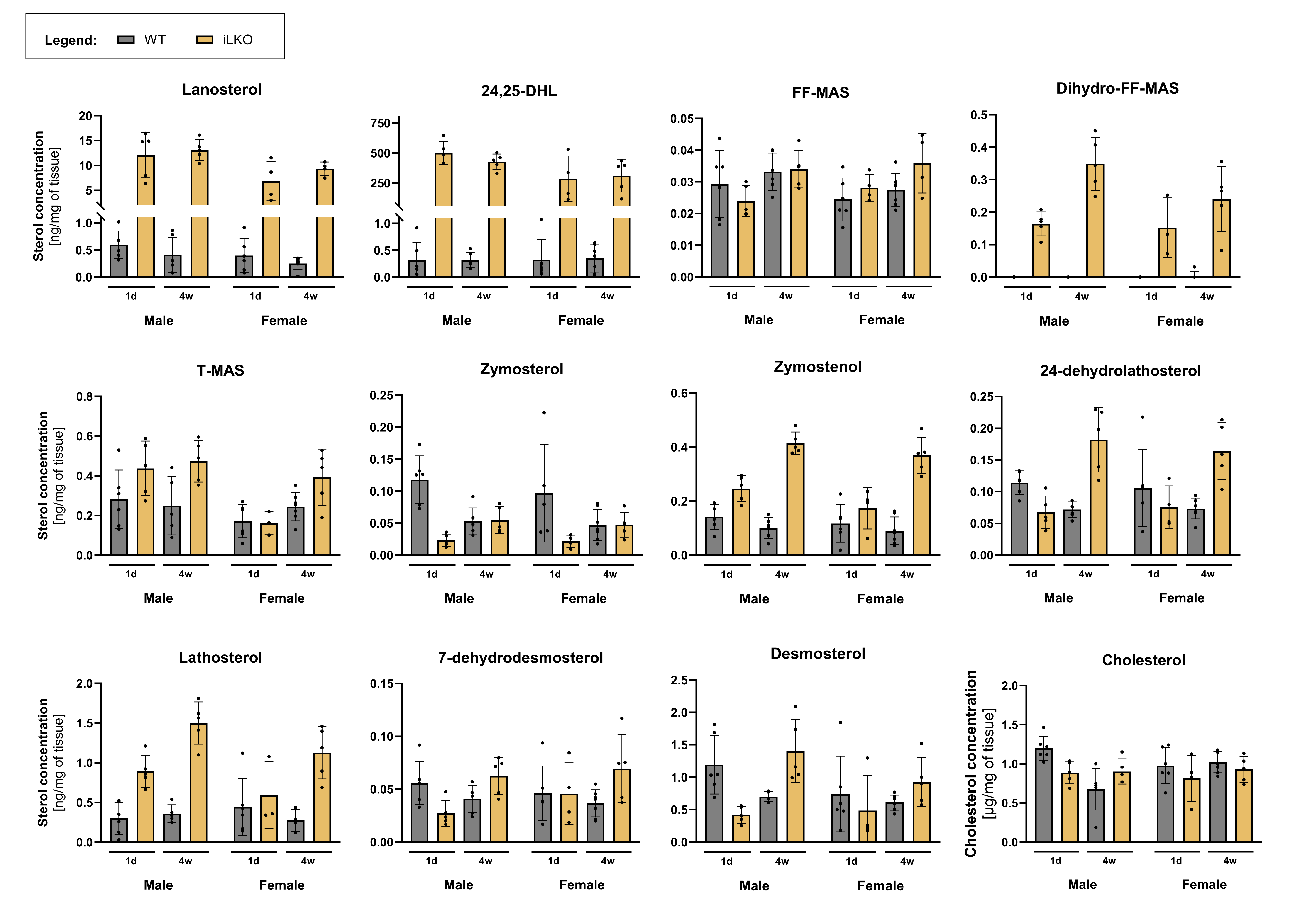

**Figure S3.** Levels of sterols extracted from liver tissue samples of iLKO and control (WT) mice. All concentrations of intermediates were calculated as ng of sterol per mg of liver tissue or in case of cholesterol µg/mg. LC-MS/MS data are represented as mean ± SD (n=4-7/group). In case of dihydro-FF-MAS, almost all WT samples exhibited levels under the quantification limit. Statistical evaluation with three-way ANOVA can be seen in Table S1. 24,25-DHL, 24,25-dihydrolanosterol.

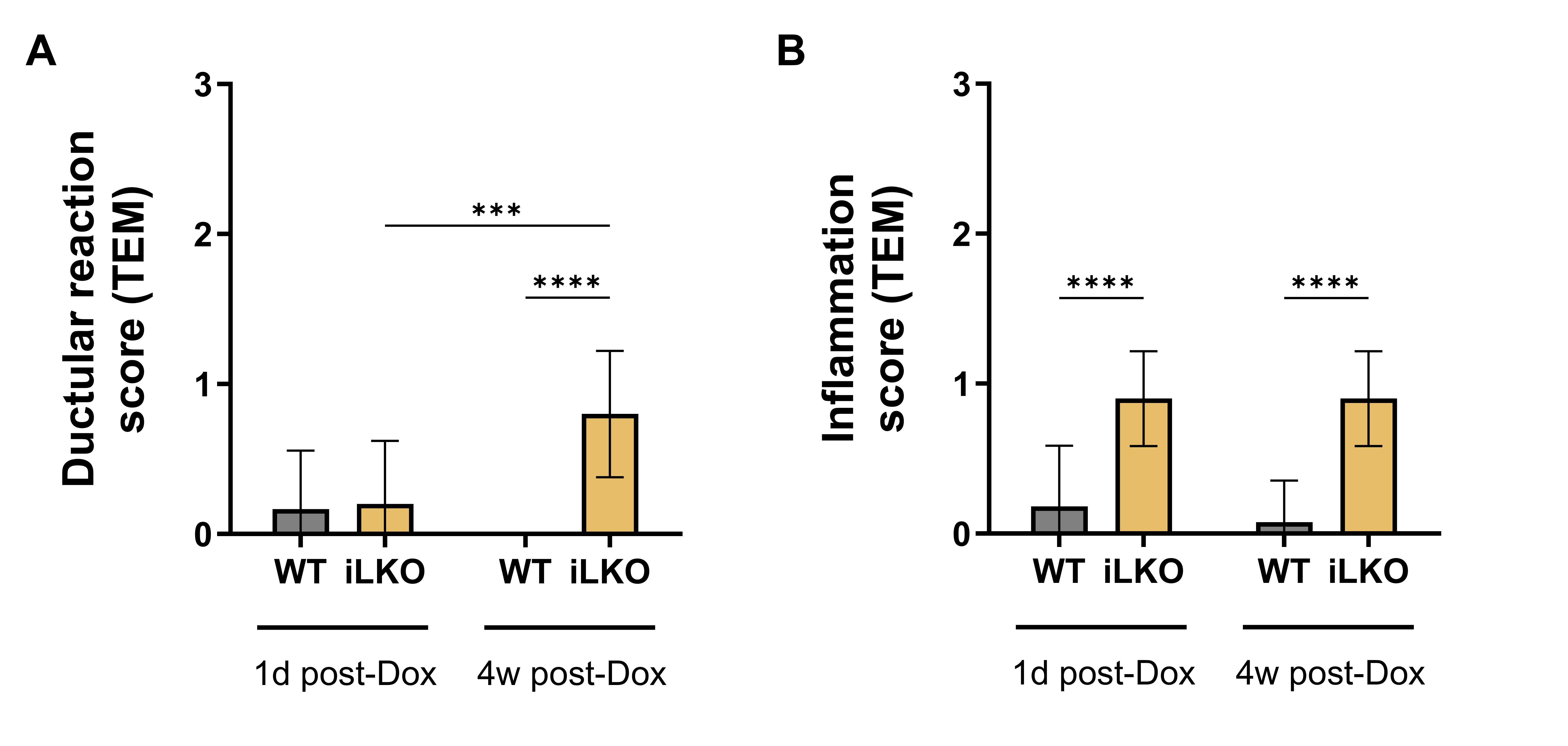

**Figure S4.** Transmission electron microscopy (TEM) semi-quantification data of (A) ductular reaction and (B) surrounding inflammation in liver samples of iLKO and WT mice, terminated 1 day or 4 weeks after last Dox application (n=10-13/group). Both processes were graded on a 0-3 scale and analyzed with two-way ANOVA and uncorrected Fisher’s test. Columns represent means ± SDs. Stat. significance is only shown for WT vs iLKO comparisons and between termination points.* p < 0.05; ** p < 0.01; *** p < 0.001; **** p < 0.0001.

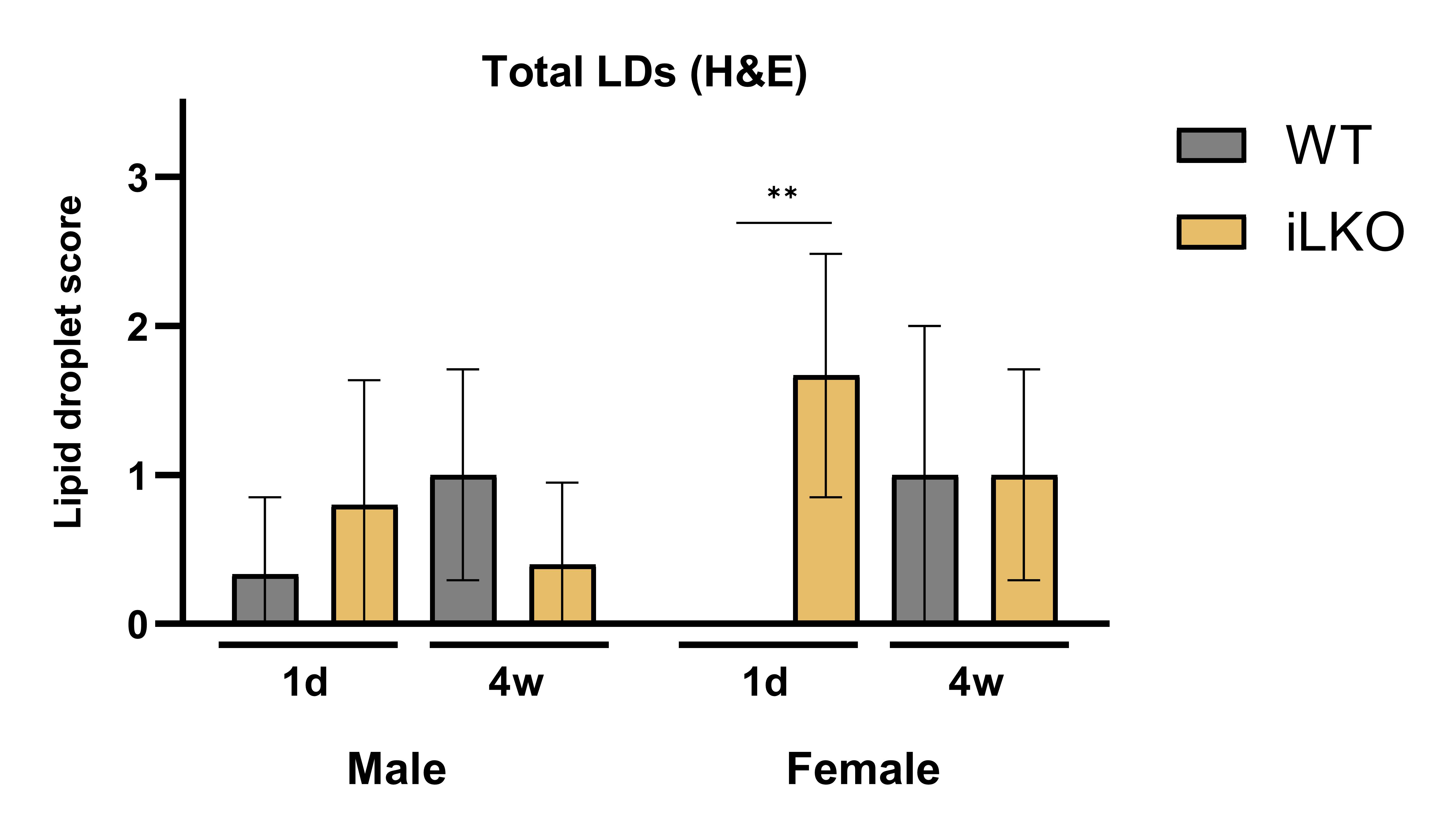

**Figure S5**. Lipid droplets (LDs) semi-quantification data obtained from Hematoxylin & Eosin (H&E) staining of wild type (WT) and iLKO liver samples. LDs presence was graded on a 0-3 scale and analyzed with three-way ANOVA and Šidak multiple comparisons test. Samples were grouped based on genotype (WT or iLKO), sex (female or male) and time of termination (1 day or 4 weeks post doxycycline (Dox) treatment). Columns represent means ± SDs (n=4-7/group); ** p < 0.01.

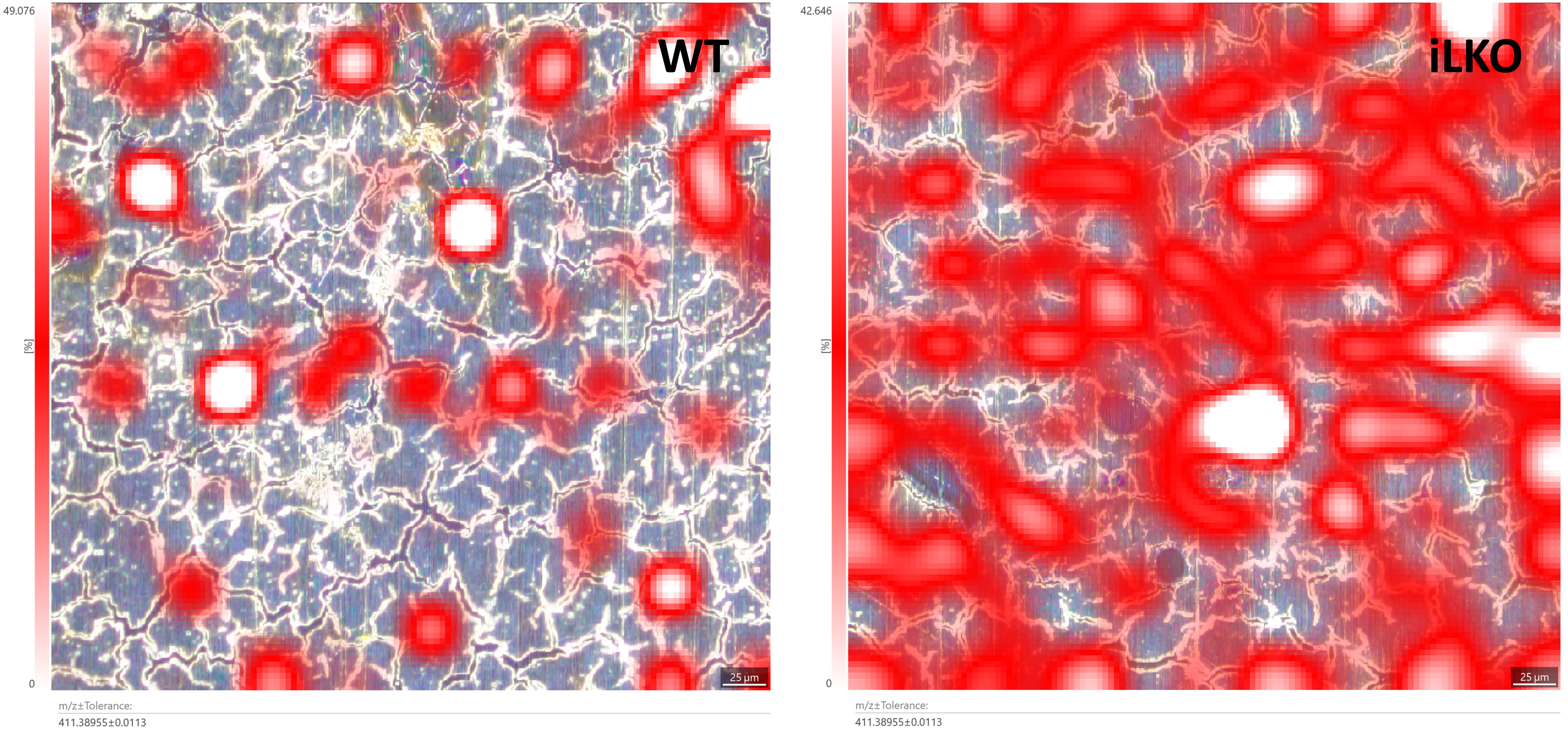

**Figure S6.** Light microscopy image of wild type (WT) (left) and iLKO (right) liver overlaid with the lateral distribution of 24,25-dihydrolanosterol (m/z value of 411.38955 ± 0.0113). Scale bar: 25 µm. Sterol distribution was evaluated with matrix-assisted laser desorption ionization–time of flight mass spectrometry imaging (MALDI-TOF MSI).

**Table S1.** List of primers for PCR analysis of transgenic mouse gDNA ear samples.

| Primer pairs | | Primer sequence (5´- 3´) | PCR cycling program | Cycles | Size amplicon (bp) |
| --- | --- | --- | --- | --- | --- |
| **Luc** | Forward | TTACAGATGCACATATCGAGG | 95 °C (3 min), 95 °C (1 min), 58 °C (1 min), 72 °C (1 min), 72 °C (7 min), 4 °C | 30 | 500 |
|  | Reverse | TAACCCAGTAGATCCAGAGG |  |  |  |
| **Cyp51 (5lox – 3lox)** | Forward | CAGACTTGATGGCAAGAGAT | 94°C (5 min), 94 °C (1 min), 59 °C (1 min), 72 °C (1 min), 72 °C (5 min), 4 °C | 35 | WT allele: 1797 Floxed: 1881 Null (KO): 187 |
|  | Reverse | TTCCGCACCTACTGTATTTT |  |  |  |
| **Cre** | Forward | TCGCTGCATTACCGGTCGATGC | 94 °C (2 min), 94 °C (1 min), 62 °C (1 min), 72 °C (1 min), 72 °C (7 min), 4 °C | 30 | 400 |
|  | Reverse | CCATGAGTGAACGAACCTGGTCG |  |  |  |
| **rtTA** | Forward | CCATGTCTAGACTGGACAAGA | 94 °C (2 min), 94 °C (1 min), 62 °C (1 min), 72 °C (1 min), 72 °C (7 min), 4 °C | 30 | 500 |
|  | Reverse | CTCCAGGCCACATATGATTAG |  |  |  |

**Table S2**. A list of health microbiological monitoring negatively detected microorganisms in mice.

Routine health microbiological

monitoring list of transgenic mice

Adenovirus FL

Adenovirus K87

Aspiculuris tetraptera

Citrobacter rodentium

Clostridium piliforme

Corynebacterium kutscheri

Cryptosporidium spp.

Ectromelia virus

Entamoeba spp.

General parvovirus (rNS-1)

Giardia spp.

Helicobacter bilis

Klebsiella oxytoca

Klebsiella pneumoniae

Minute virus of mice

Mouse hepatitis virus

Mouse parvovirus (rVP2)

Mouse rotavirus / EDIM

Mycoplasma pulmonis

Myobia musculi / Radfordia sp.

Other ectoparasites

Pneumonia virus of mice

Pseudomonas aeruginosa

Reovirus type 3

Salmonela spp.

Sendai virus

Spironucleus spp.

Staphylococcus aureus

Streptobacillus moniformis

Streptococci beta-haemolytic Group A

Streptococci beta-haemolytic Group B

Streptococci beta-haemolytic Group C

Streptococci beta-haemolytic Group G

Streptococcus pneumoniae

Syphacia obvelata

Theiler's encephalomyelitis virus (GD VII)

**Table S3.** List of buffers and reagents using in primary hepatocyte isolation procedure.

| Buffer/reagent | | Components and preparation |
| --- | --- | --- |
| **Hanks buffer** | | NaCl (136.3 mM), KCl (5.4 mM), MgSO_4_ x 7 H_2_O (0.4 mM), MgCl_2_ x 6 H_2_O (0.5 mM), Na_2_HPO_4_ x 2 H_2_O (0.34 mM), KH_2_PO_4_ (0.44 mM), CaCl_2_ (0.13 mM), HEPES (2 mM), EDTA (1.3 mM). Components were dissolved in 1 l of sterile double deionized water; pH 7.4-7.5, autoclaved and stored at 4 °C. |
|  | Hanks buffer^+/+^ | Hanks buffer w/o EDTA. |
|  | Hanks buffer^+/+^ + EDTA | Hanks buffer with all components. |
|  | Hanks buffer^-/-^ | Hanks buffer w/o EDTA, Ca, Mg. |
| **Isolation medium (IM)** | | KCl (5.36 mM), MgSO4 x 7 H2O (0.29 mM), Na2HPO4 x 2 H2O (0.79 mM), Na2HPO4 x 2 H2O (0.15 mM), HEPES (10 mM), NaCl (145 mM). Components were dissolved in 1 l of sterile double deionized water; pH 7.4-7.5, autoclaved and stored at 4 °C. |
|  | Isolation medium (IMS) | 2.5 ml 200mM CaCl_2_ and 5 ml 10% Glucose were dissolved in 0.5 l of IM. |
| **Perfusion buffer (PM)** | | KCl (5.36 mM), MgSO_4_ x 7 H_2_O (0.77 mM), MgCl_2_ x 6 H_2_O (0.93 mM), Na_2_HPO_4_ x 2 H_2_O (0.34 mM), KH_2_PO_4_ (0.44 mM), HEPES (10 mM), NaCl (145 mM), Glucose (0.2%), Penicillin (1653U/mg), Streptomycin (757U/mG), BSA Albumin Frakt V 0.2 %. Components were dissolved in 1 l of sterile double deionized water; pH 7.4-7.5, sterile filtrated and stored in 50 ml aliquots in the freezer at -20 °C. |
| **Isolation perfusion buffer (PPM)** | | KCl (5.36 mM), MgSO_4_ x 7 H_2_O (0.77 mM), MgCl_2_ x 6 H_2_O (0.93 mM), Na_2_HPO_4_ x 2 H_2_O (0.34 mM), KH_2_PO_4_ (0.44 mM), EGTA (0.2 mM), HEPES (10 mM), NaCl (145 mM). Components were dissolved in 1 l of sterile double deionized water; pH 7.4-7.5, autoclaved and stored at 4 °C. |
| **Collagenase** | | Collagenase type I, Sigma-Aldrich, St. Louis, MO, USA. The dedicated amount of collagenase was dissolved in 10 ml of Hanks buffer^-/-^ buffer and sterile-filtrated freshly before the procedure. |
| **200 mM CaCl_2_** | |  |
| **10% Glucose** | |  |

**Table S4.** List of primers used in RT-qPCR analyses.

| **Gene Symbol** | **Official gene name** | **Forward primer sequence (5´- 3´)** | **Reverse primer sequence (5´- 3´)** |
| --- | --- | --- | --- |
| ***Cyp51*** | Cytochrome P450 family 51 | ACGCTGCCTGGCTATTGC | TTGATCTCTCGATGGGCTCTATC |
| ***Utp6*** | Small subunit processome component | TTTCGGTTGAGTTTTTCAGGA | CCCTCAGGTTTACCATCTTGC |
| ***Hmbs*** | hydroxymethylbilane synthase | TCCCTGAAGGATGTGCCTA | AAGGGTTTTCCCGTTTGC |
| ***Actb*** | Actin beta | CTTCCTCCCTGGAGAAGAGC | ATGCCACAGGATTCCATACC |

**Table S5.** MS detection conditions of the postsqualene cholesterol synthesis sterol intermediates in primary hepatocytes and liver tissue of iLKO and control mice.

| Trivial Name | Systematic name | MRM | Molar mass |
| --- | --- | --- | --- |
| Lanosterol | lanosta-8,24-dien-3β-ol | 409/191 | 426.72 |
| Dihydrolanosterol | 24,25-dihydrolanosterol | 411/191 | 428.73 |
| FF-MAS * | 14-demethyl-14-dehydrolanosterol | 393/214 | 410.68 |
| T-MAS * | 4,4-dimethylcholest-8(9),24-dien-3β-ol | 395/243 | 412.69 |
| Desmosterol | cholest-5,24-dien-3β-ol | 367/215 | 384.64 |
| 24-dehydrolathosterol | 5α-cholesta-7,24-dien-3β-ol | 367/215 | 384.64 |
| Lathosterol-d7 | 5α-Cholest-7-en-3β-ol(25,2626,26,27,27,27-d7) | 376/215 | 393.7 |
| Zymosterol* | 5α-cholesta-8,24-dien-3β-ol | 367/215 | 384.64 |
| Zymostenol | 5α-cholest-8-en-3β-ol | 369/215 | 386.65 |
| Cholesterol | cholest-5-en-3β-ol | 369/215 | 386.7 |
| Dihydro-FF-MAS ** | 4,4-dimethyl-5α-cholesta-8,14-dien-3β-ol | 395/123 | 412.37 |
| Lathosterol ** | cholest-7-en-3β-ol | 369/215 | 383.35 |
| 7-dehydrodesmosterol ** | cholest-5,7,24-trien-3β-ol | 365/199 | 382.32 |

*Sterol concentration was too low to be quantified in primary hepatocytes samples.

**Sterol additionally added to the method and measured only in tissue samples.

**Table S6.** Statistical analysis of measured sterol levels extracted from liver tissue samples of iLKO and control (WT) mice. Subjects were grouped based on genotype, sex and time of termination (denoted ‘Age’) (n=4-7/group). All concentrations of intermediates were calculated as ng of sterol per mg of liver tissue or in case of cholesterol µg/mg. Ordinary three-way ANOVA was applied; ns, not stat. significant; * p < 0.05; ** p < 0.01; *** p < 0.001; **** p < 0.0001.

|  | **Lanosterol** | **% of total variation** | **P value** | |  |  | **Zymostenol** | **% of total variation** | **P value** | |
| --- | --- | --- | --- | --- | --- | --- | --- | --- | --- | --- |
| Source of variation | Age | 0.4839 | 0.2332 | ns |  | Source of variation | Age | 8.308 | 0.0001 | *** |
|  | Genotype | 75.91 | <0.0001 | **** |  |  | Genotype | 54.36 | <0.0001 | **** |
|  | Sex | 4.241 | 0.001 | *** |  |  | Sex | 2.253 | 0.029 | * |
|  | Age x Genotype | 0.7077 | 0.1512 | ns |  |  | Age x Genotype | 17.78 | <0.0001 | **** |
|  | Age x Sex | 0.1087 | 0.569 | ns |  |  | Age x Sex | 0.1573 | 0.5516 | ns |
|  | Genotype x Sex | 3.618 | 0.0021 | ** |  |  | Genotype x Sex | 0.6533 | 0.2285 | ns |
|  | Age x Genotype x Sex | 0.09734 | 0.5898 | ns |  |  | Age x Genotype x Sex | 0.01363 | 0.8605 | ns |
|  | **24,25-dihydrolanosterol** | **% of total variation** | **P value** | |  |  | **24-dehydrolathosterol** | **% of total variation** | **P value** | |
| Source of variation | Age | 0.07867 | 0.6275 | ns |  | Source of variation | Age | 9.289 | 0.0063 | ** |
|  | Genotype | 78.39 | <0.0001 | **** |  |  | Genotype | 8.662 | 0.0081 | ** |
|  | Sex | 3.666 | 0.002 | ** |  |  | Sex | 0.1744 | 0.6931 | ns |
|  | Age x Genotype | 0.07888 | 0.6271 | ns |  |  | Age x Genotype | 43.41 | <0.0001 | **** |
|  | Age x Sex | 0.3326 | 0.321 | ns |  |  | Age x Sex | 0.1486 | 0.7156 | ns |
|  | Genotype x Sex | 3.667 | 0.002 | ** |  |  | Genotype x Sex | 0.002813 | 0.96 | ns |
|  | Age x Genotype x Sex | 0.3324 | 0.3211 | ns |  |  | Age x Genotype x Sex | 0.7608 | 0.4115 | ns |
|  | **FF-MAS** | **% of total variation** | **P value** | |  |  | **Lathosterol** | **% of total variation** | **P value** | |
| Source of variation | Age | 18.17 | 0.0033 | ** |  | Source of variation | Age | 2.854 | 0.0634 | ns |
|  | Genotype | 2.553 | 0.2464 | ns |  |  | Genotype | 56.6 | <0.0001 | **** |
|  | Sex | 0.08363 | 0.8323 | ns |  |  | Sex | 0.7035 | 0.3479 | ns |
|  | Age x Genotype | 4.491 | 0.1268 | ns |  |  | Age x Genotype | 6.941 | 0.005 | ** |
|  | Age x Sex | 0.005859 | 0.9553 | ns |  |  | Age x Sex | 1.756 | 0.1416 | ns |
|  | Genotype x Sex | 9.024 | 0.0331 | * |  |  | Genotype x Sex | 1.47 | 0.1776 | ns |
|  | Age x Genotype x Sex | 0.02391 | 0.9098 | ns |  |  | Age x Genotype x Sex | 0.08306 | 0.7457 | ns |
|  | **Dihydro-FF-MAS** | **% of total variation** | **P value** | |  |  | **7-dehydrodesmosterol** | **% of total variation** | **P value** | |
| Source of variation | Age | 8.54 | <0.0001 | **** |  | Source of variation | Age | 3.516 | 0.1837 | ns |
|  | Genotype | 63.87 | <0.0001 | **** |  |  | Genotype | 1.863 | 0.3305 | ns |
|  | Sex | 2.043 | 0.0269 | * |  |  | Sex | 0.3686 | 0.6634 | ns |
|  | Age x Genotype | 8.062 | <0.0001 | **** |  |  | Age x Genotype | 20.34 | 0.0024 | ** |
|  | Age x Sex | 0.2529 | 0.4223 | ns |  |  | Age x Sex | 0.1245 | 0.8001 | ns |
|  | Genotype x Sex | 2.287 | 0.0197 | * |  |  | Genotype x Sex | 4.56 | 0.1315 | ns |
|  | Age x Genotype x Sex | 0.3431 | 0.3508 | ns |  |  | Age x Genotype x Sex | 0.8745 | 0.5034 | ns |
|  | **T-MAS** | **% of total variation** | **P value** | |  |  | **Desmosterol** | **% of total variation** | **P value** | |
| Source of variation | Age | 5.034 | 0.067 | ns |  | Source of variation | Age | 3.444 | 0.1435 | ns |
|  | Genotype | 18.99 | 0.0008 | *** |  |  | Genotype | 0.0434 | 0.8676 | ns |
|  | Sex | 12.78 | 0.0048 | ** |  |  | Sex | 5.117 | 0.0766 | ns |
|  | Age x Genotype | 4.056 | 0.0986 | ns |  |  | Age x Genotype | 30.67 | <0.0001 | **** |
|  | Age x Sex | 6.706 | 0.0359 | * |  |  | Age x Sex | 0.06217 | 0.8419 | ns |
|  | Genotype x Sex | 4.576 | 0.0801 | ns |  |  | Genotype x Sex | 0.01646 | 0.9182 | ns |
|  | Age x Genotype x Sex | 0.2521 | 0.6749 | ns |  |  | Age x Genotype x Sex | 6.597 | 0.0457 | * |
|  | **Zymosterol** | **% of total variation** | **P value** | |  |  | **Cholesterol** | **% of total variation** | **P value** | |
| Source of variation | Age | 1.574 | 0.3127 | ns |  | Source of variation | Age | 3.529 | 0.1523 | ns |
|  | Genotype | 19.1 | 0.001 | ** |  |  | Genotype | 3.218 | 0.1711 | ns |
|  | Sex | 1.893 | 0.2689 | ns |  |  | Sex | 0.1562 | 0.7601 | ns |
|  | Age x Genotype | 20.34 | 0.0008 | *** |  |  | Age x Genotype | 10.5 | 0.0162 | * |
|  | Age x Sex | 0.4166 | 0.6015 | ns |  |  | Age x Sex | 12.69 | 0.0087 | ** |
|  | Genotype x Sex | 0.7501 | 0.4841 | ns |  |  | Genotype x Sex | 0.7966 | 0.4916 | ns |
|  | Age x Genotype x Sex | 0.9333 | 0.4355 | ns |  |  | Age x Genotype x Sex | 6.169 | 0.0611 | ns |

**Table S7.** The size of 20 crystal-like structures in different iLKO liver samples by transmission electron microscopy (TEM).

|  | Width (nm) | Length (nm) |
| --- | --- | --- |
| 1 | 86 | 624 |
| 2 | 50 | 54 |
| 3 | 50 | 43 |
| 4 | 92 | 573 |
| 5 | 70 | 1820 |
| 6 | 140 | 57 |
| 7 | 108 | 1150 |
| 8 | 48 | 71 |
| 9 | 122 | 961 |
| 10 | 144 | 785 |
| 11 | 150 | 3900 |
| 12 | 218 | 875 |
| 13 | 75 | 1625 |
| 14 | 66 | 1693 |
| 15 | 133 | 1320 |
| 16 | 80 | 1200 |
| 17 | 280 | 1200 |
| 18 | 32 | 504 |
| 19 | 56 | 672 |
| 20 | 112 | 1120 |
| Median (95% CI) | 89 (66-133) | 918 (573-1200) |
